# AM-DMF-SCP-pro: An Ultrafast and Streamlined Nanolitre-Scale Lab-on-a-Chip Platform for Single-Cell Proteomics and Tumor Microenvironment Profiling

**DOI:** 10.64898/2026.04.29.721532

**Authors:** Kai Jin, Zhicheng Yang, Anqi Ding, Rong Wang, Yifan Ma, Yuqiu Wang, Minchu Tang, Yong Wu, Qian Liu, Jiajian Ji, Chunyu Chang, Siyi Hu, Wenfei Dong, Jian Ding, Yi Chen, Hanbin Ma, Hu Zhou

## Abstract

Single-cell proteomics (SCP) remains constrained by slow sample preparation and background interference in low-input, multistep workflows, limiting its broader practical use in complex biological samples. Here we report AM-DMF-SCP-pro, a streamlined active-matrix digital microfluidic workflow that enables rapid on-chip lysis and digestion in 6 nL droplets within a closed nanolitre-confined microenvironment. AM-DMF-SCP-pro reduces total sample-preparation time to as little as 30 min, including a 15 min digestion step, while maintaining quantitative reproducibility and digestion performance. Systematic optimization of mass spectrometry acquisition and nanolitre-scale digestion conditions enables deeper, more sensitive and lower-background proteome profiling from ultra-low peptide inputs. In murine tumor samples, AM-DMF-SCP-pro reliably resolves immune and tumor compartments, identifies treatment-associated tumor cell states and enables quantitative analysis of neutrophil-associated proteomic signatures. These results establish AM-DMF-SCP-pro as a practical workflow for rapid, low-background single-cell proteome profiling in complex biological samples.

## Introduction

Elucidating the molecular heterogeneity of complex biological systems requires precision tools capable of resolving functional states at the single-cell level^1,2^. While single-cell RNA sequencing (scRNA-seq) has provided extensive atlases of cell types, cellular function is ultimately executed by proteins, whose abundances often correlate poorly with their corresponding mRNAs due to post-transcriptional regulation, degradation, and transport^3,4^. Consequently, mass spectrometry (MS)-based single-cell proteomics has emerged as an essential complement to transcriptomics, offering direct measurements of enzymatic activities, signalling pathways, and protein isoforms that define cellular phenotypes^5–8^. However, unlike nucleic acids, proteins cannot be amplified, making SCP inherently susceptible to sample loss during preparation, thereby severely limiting analytical sensitivity and proteome coverage^9,10^.

A persistent challenge in SCP workflows is the physical handling of picogram-level protein inputs^11^. Traditional preparation methods using microliter volumes in standard tubes or well plates suffer from high surface-to-volume ratios, leading to substantial adsorption of hydrophobic peptides to the vessel walls. To mitigate this, recent container surface functionalization strategies and miniaturized platforms such as OAD^12^, nanoPOTS^13^, nPOP^14^, Mad-CASP^15^, and related systems^16–21^ have reduced protein adsorption to surfaces and sample loss, thereby improving analytical sensitivity. However, many platforms still rely on specialized liquid-handling systems, such as CellenONE^22,23^, HP D100^24^ or custom-built dispensers^25,26^, and are often implemented on open substrates, including well plates or glass surfaces. These workflows frequently involve multistep reagent additions, increasing operational variability as well as the risk of exogenous contamination and evaporation, and often require digestion times of several hours, which prolong sample preparation and limit throughput. Together, these limitations continue to constrain the broader and more scalable application of SCP in the life sciences^8,27,28^.

Digital microfluidics (DMF), which manipulates discrete droplets on electrode array via electrowetting-on-dielectric (EWOD), offers a closed, programmable and automated alternative for proteomic sample preparation^29,30^. Although active-matrix digital microfluidics (AM-DMF) has established the feasibility of on-chip sample preparation for single-cell proteomics^31,32^, previous AM-DMF workflows still relied on multistep processing and hour-scale digestion, which required larger on-chip areas and prolonged sample-preparation time, thereby limiting parallelization. However, the confined nanolitre-scale microenvironment of DMF provides elevated effective analyte concentrations and shorter diffusion and mixing distances, suggesting that on-chip SCP workflows can be substantially streamlined through fewer reaction steps, smaller reaction volumes and shorter sample-preparation times.

Here, we present AM-DMF-SCP-pro, an ultrafast and streamlined nanolitre-scale single-cell proteomics platform based on active-matrix digital microfluidics. AM-DMF-SCP-pro performs on-chip lysis and digestion in 6 nL droplets within a closed nanolitre-confined microenvironment. AM-DMF-SCP-pro reduces total sample-preparation time to 30 min, including a 15 min digestion step, which, to our knowledge, represents the shortest total sample-preparation time and fastest digestion reported for SCP. We systematically optimized key components of the workflow, including multi-droplet routing, MS acquisition parameters and nanolitre-scale lysis–digestion conditions, to improve processing efficiency, proteomic sensitivity and analytical robustness. We further benchmarked AM-DMF-SCP-pro against plate-based SCP workflows, demonstrating reduced background interference and improved signal-to-noise in nanolitre-scale single-cell proteomics. To further investigate cellular heterogeneity and functional states in the tumor microenvironment, we apply the platform to murine tumors and examine treatment-associated changes in both tumor and immune compartments. In particular, we use this framework to characterize neutrophil-associated proteomic responses to anti-angiogenic therapy, highlighting the value of single-cell proteomics for resolving functional features that are not readily captured by transcriptomic approaches alone.

## Results

### Development and Integration of an Automated, Nanolitre-Scale Digital Microfluidic Platform for Ultrafast Single-Cell Proteomics

We engineered the AM-DMF-SCP-pro system to seamlessly integrate the entire proteomic pipeline-from the initial isolation of single cells to peptide collection-within a single microfluidic chip (Figures 1A-C). Unlike traditional SCP workflows that involve a series of discrete, time-consuming steps such as lysis, reduction, alkylation, and digestion, often spanning 3.5 to 16 hours^17,22,33,34^, our method combines the nanolitre-confined microenvironment of DMF with a streamlined lysis-digestion workflow, reducing total sample-preparation time to 0.5-0.75 h (Figure 1D). Digestion was shortened from at least 2 h in previous workflows to 15 min, representing an approximately eightfold reduction and laying the technical foundation for higher daily throughput. AI-guided path planning enables precise and parallel manipulation of single-cell and reagent droplets on the DMF chip, including droplet routing, merging, and mixing (Figure 1E). These operations are performed within a DMF cartridge that provides a confined nanolitre-scale microenvironment for single-cell sample preparation (Figure 1F). To visualize a key operation in the workflow, a schematic top view shows the merger of a single-cell droplet with reagent droplets within the cartridge (Figure 1G). Through meticulous optimization of the on-chip protocol, the entire process duration has been reduced to as little as 30 minutes This significant acceleration leverages the physics of nanolitre reactors, which minimize diffusion distances and effectively concentrate reagents, thereby dramatically enhancing reaction effect and overall system efficiency.

**Figure 1.**
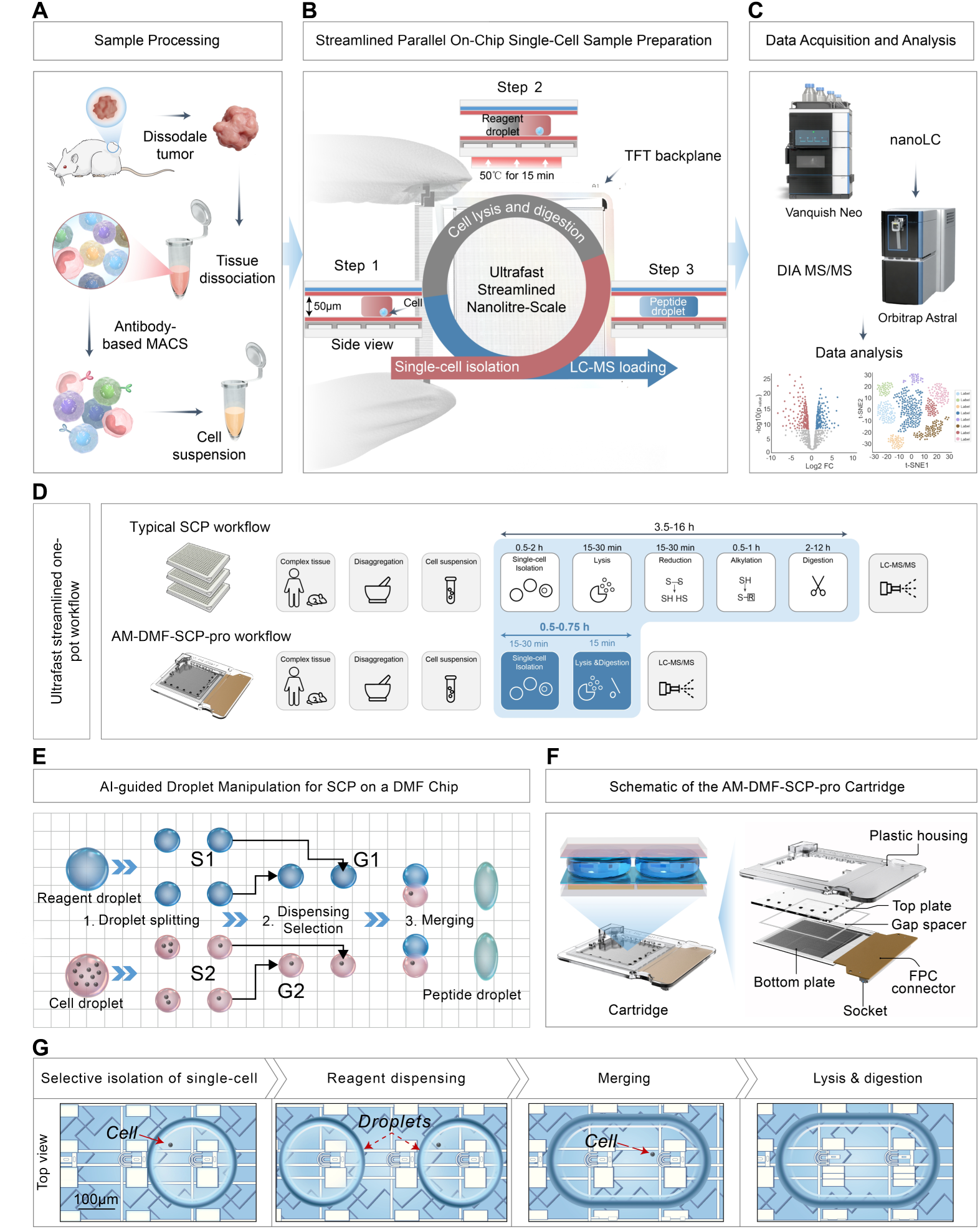
Overview of the ultrafast and streamlined single-cell proteomics workflow enabled by a digital microfluidic lab-on-chip. (A) Processing from tissue specimens to single-cell suspensions. (B) Streamlined DMF lab-on-chip workflow for parallel single-cell sample preparation, including on-chip single-cell isolation, lysis and digestion, prepared samples are transferred directly to LC-MS/MS. (C) Single-cell proteomic data acquisition and analysis. (D) Comparison between the proposed ultrafast, automated lab-on-a-chip SCP workflow and a typical SCP workflow. Our workflow completes single-cell lysis and digestion within 15 minutes, 8 times faster than typical protocols. (E) Schematic of AI-guided droplet manipulation within the DMF cartridge, showing automated and parallel handling of single-cell and reagent droplets. (F) Exploded view of the DMF cartridge, which provides a confined nanolitre-scale microenvironment for single-cell sample preparation. (G) A schematic top view shows the merger between a single-cell droplet and reagent droplets, clarifying this key step in the workflow.

### Enhanced Safe-Interval Path Planning and a Streamlined DMF Workflow Reduce Reaction Steps and Processing Time for Higher Daily Throughput

Inspired by multi-agent path-planning concepts from artificial intelligence and robotics^35,36^, we sought to improve coordinated droplet manipulation for automated single-cell sample preparation on a digital microfluidic platform. Building on our previously reported SIPP-based droplet-routing approach^37,38^, we implemented an enhanced safe-interval path planning framework to address the increasing computational cost of routing under dense multi-droplet conditions (Figure 2A). Across benchmark scenarios containing 10-100 single-cell droplets, the enhanced SIPP implementation consistently completed route planning in under 1 s, whereas the earlier SIPP-based implementation required progressively longer computation as droplet number increased and reached ∼24 s at the largest tested scale (Figure 2B). Despite the substantial reduction in planning time, the overall makespan remained comparable between the two methods across the same range of droplet numbers (Figure 2C), indicating improved computational efficiency without an evident loss in routing compactness. Representative top-view experimental frames and corresponding software screenshots showed close agreement between planned and executed droplet trajectories during multi-droplet routing on chip (Figure 2D). Building on this routing performance, we implemented an automated workflow for single-cell sample preparation in the DMF cartridge and incorporated a streamlined lysis-digestion strategy to further reduce handling complexity and processing time (Figure 2E, Figure S1, Supplementary Movie 1). Compared with the AM-DMF-SCP workflow reported in 2024, the present workflow requires a smaller reaction volume and fewer reaction steps, while also reducing total sample-preparation time (Figure 2F, Figure S2). Together, these results demonstrate that enhanced SIPP enables scalable and reliable on-chip multi-droplet routing, while the streamlined DMF workflow reduces reaction volume, simplifies sample processing, and shortens processing time, thereby providing a practical basis for higher-throughput single-cell sample preparation.

**Figure 2.**
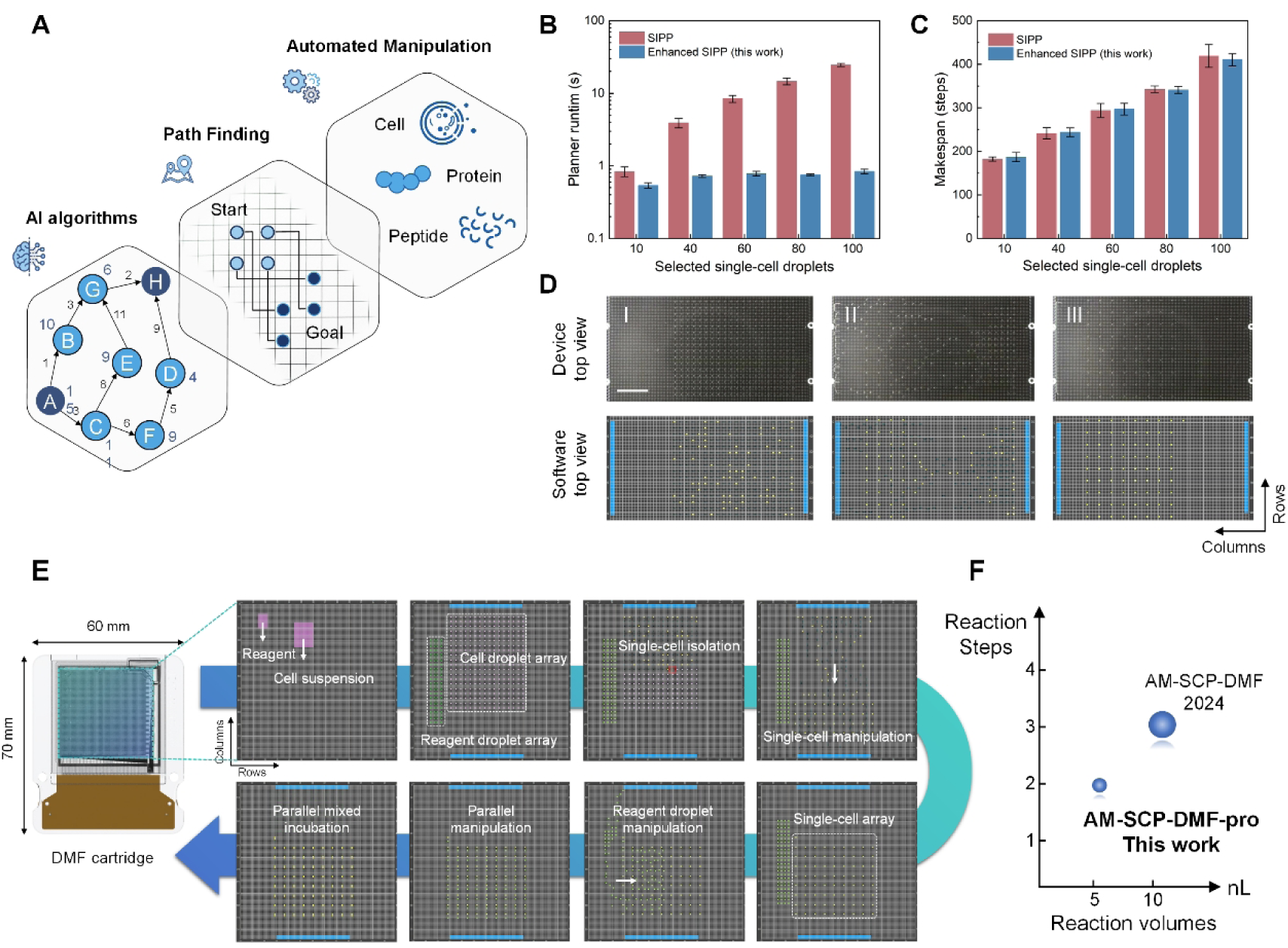
Enhanced safe-interval path planning and streamlined workflow for automated single-cell sample preparation on the digital microfluidic platform. A. Schematic illustration of the routing framework, showing how AI-based path planning enables single-cell droplet manipulation on the digital microfluidic platform for automated sample preparation. B. Planning time of conventional SIPP and enhanced SIPP for 10-100 single-cell droplets. C. Makespan (steps) of the two methods across the same test conditions. D. Representative device and software top views during multi-droplet routing. Shown are three experimental top-view frames of on-chip droplet actuation and three corresponding software top-view screenshots, illustrating agreement between planned and executed droplet trajectories. The scale bar is 5 mm. E. Schematic of the overall workflow for single-cell sample preparation in the DMF cartridge (60 mm × 70 mm). F. Compared with AM-SCP-DMF 2024, this work requires a smaller reaction volume and fewer reaction steps.

### Systematic optimization of MS acquisition parameters and a streamlined lysis-digestion workflow enhance sensitivity in AM-DMF-based single-cell proteomics

In single-cell proteomics, the extremely low sample input makes MS acquisition parameters a critical determinant of proteome coverage and quantitative stability. We therefore systematically optimized MS acquisition parameters using ultra-low-input peptide samples (50 pg HEK-293T digest), including DIA isolation window width and FAIMS compensation voltage^39^ (Figure 3A, B and Figure S3A, B). We found that a 20 Th isolation window combined with a FAIMS CV of −48 V provided optimal performance. Under these conditions, up to 5,916 protein groups (PGs) were identified from 250 pg input (Figure S3C, D). In addition, accurate ratio recovery was achieved in mixed-species samples, indicating a balanced performance between proteome depth and quantitative accuracy (Figure S4).

**Figure 3.**
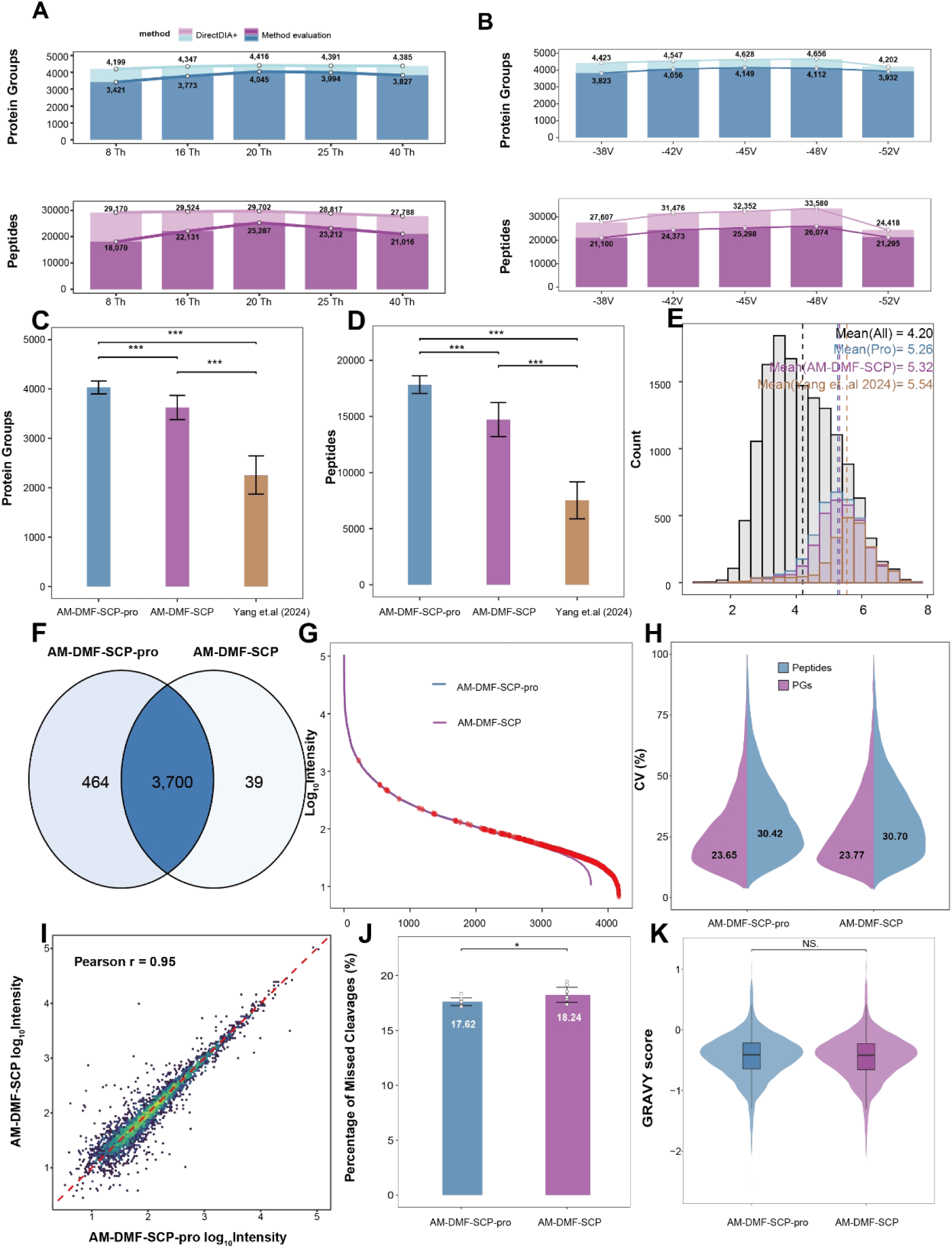
Streamlined workflow enhances sensitivity while preserving quantitative robustness in AM-DMF-based single-cell proteomics. (A) Effect of isolation window on protein identification. (B) Effect of compensation voltage on protein identification. (C) Comparison of protein group coverage among AM-DMF-SCP-pro, AM-DMF-SCP, and the previously reported data from Yang et al. (2024). (D) Comparison of peptide coverage for each condition. (E) Distribution of quantified PGs per cell, with dashed lines indicating mean values for each condition. (F) Overlap of identified PGs between the two workflows. (G) Comparison of ranked protein group intensities between the two workflows. PGs unique to AM-DMF-SCP-pro are highlighted in red. (H) Distribution of the coefficients of variation (CV) of protein intensities for the two workflows. (I) Pairwise correlation of protein intensities between the two workflows, showing overall quantitative agreement. (J) Comparison of missed-cleavage percentages between the two workflows. (K) Distribution of GRAVY scores for proteins identified in the two workflows. AM-DMF-SCP-pro and AM-DMF-SCP were evaluated under the optimized mass spectrometry conditions used in this study, whereas Yang et al. (2024) refers to AM-DMF-SCP data acquired under earlier mass spectrometry conditions.

Under these optimized MS conditions, we further evaluated the performance of the streamlined lysis-digestion strategy of AM-DMF-SCP-pro in true single-cell proteomics experiments using HeLa cells. Compared with AM-DMF-SCP and previously reported data^31^, AM-DMF-SCP-pro achieved improved protein-group and peptide coverage (Figure 3C,D). Analysis of protein copy number distributions^40^ revealed a clear extension toward lower copy-number proteins in the AM-DMF-SCP-pro workflow (Figure 3E), demonstrating enhanced sensitivity for low-abundance proteins. Overlap analysis further showed that most proteins were consistently identified in both workflows, while 425 additional proteins were uniquely detected in the AM-DMF-SCP-pro condition (Figure 3F), supporting its improved sensitivity for lower-abundance proteins. Consistently, the rank-intensity plot showed similar overall abundance distributions between the two workflows, whereas PGs uniquely identified in AM-DMF-SCP-pro were concentrated in the lower-intensity range (Figure 3G, red). Despite this improvement in depth and sensitivity, quantitative consistency was well preserved. The distributions of coefficients of variation (CVs) for quantified PGs and peptides were comparable between the AM-DMF-SCP-pro and AM-DMF-SCP workflows (Figure 3H), indicating that the streamlined workflow preserves quantitative reproducibility. Median CVs for PGs were 23.65% and 23.77%, respectively, whereas median CVs for peptides were 30.42% and 30.70%. Consistently, strong correlations were observed between the two workflows (Pearson r = 0.95) (Figure 3I). Digestion quality also remained comparable, with similar missed-cleavage percentages in AM-DMF-SCP-pro and AM-DMF-SCP (17.62% and 18.24%, respectively; Figure 3J). In addition, similar GRAVY score distributions were observed between the two workflows (Figure 3K), indicating that the streamlined workflow does not introduce an obvious hydrophobicity bias or preferential loss of hydrophobic proteins. Together, these results demonstrate that systematic optimization of MS acquisition parameters and the streamlined lysis-digestion workflow enhance proteome depth and sensitivity in AM-DMF-based single-cell proteomics without compromising quantitative reproducibility or digestion quality.

### Nanolitre-scale digestion kinetics enable rapid and efficient single-cell proteome preparation

In single-cell proteomics, enzymatic digestion is a major time-consuming step that constrains overall throughput. Given the confined nanolitre-scale microenvironment of AM-DMF-SCP-pro, we systematically optimized and quantitatively evaluated digestion conditions to assess whether rapid digestion could be achieved without loss of performance. We first monitored single-cell lysis dynamics using microscopy. Complete cell lysis was achieved within approximately 40 s after merging the lysis reagent with the cell-containing droplet (Figure 4A, Supplementary Movie 2), indicating that cell disruption is rapid in the nanolitre system and that downstream digestion becomes the primary limiting step. Given that nanolitre-scale reactions exhibit higher effective substrate concentrations and shorter diffusion distances compared to conventional microliter-scale systems, we hypothesized that enzymatic reaction kinetics would be significantly accelerated (Figure S5A). To test this, we systematically evaluated the effects of enzyme concentration (20 ng/µL and 120 ng/µL) and digestion time (5-120 min) on single HeLa cell (Figure 4B). Missed cleavage rates decreased with increasing digestion time under both conditions. Notably, at high enzyme concentration, a digestion time of 15 min resulted in a missed cleavage rate of 19.34%, which was comparable to that obtained at low enzyme concentration after 120 min (18.91%) (Figure 4C), indicating that increased enzyme concentration markedly accelerates digestion. Consistently, trypsin autodigestion signals increased with enzyme concentration (Figure S5B).

**Figure 4.**
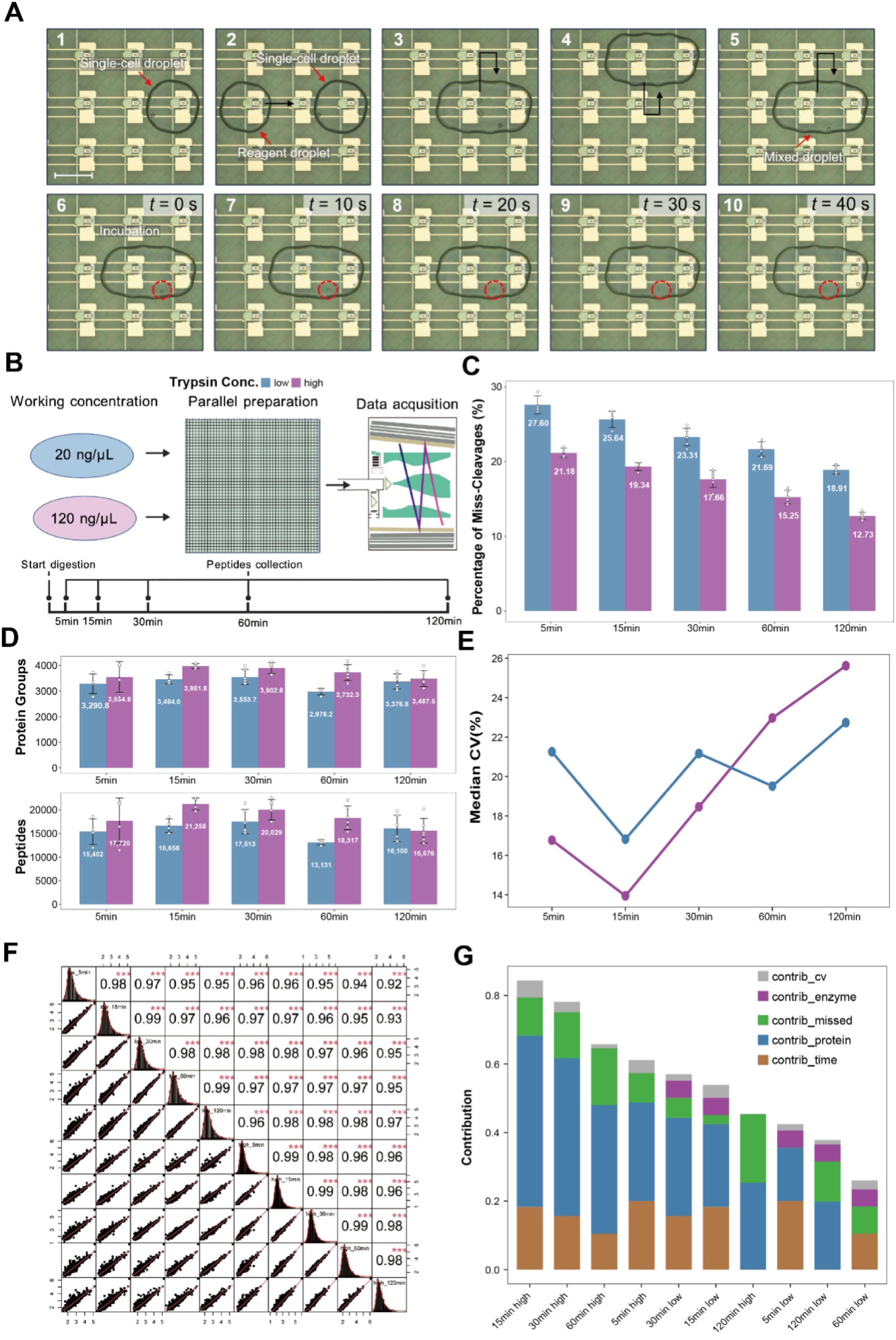
Optimization of enzymatic digestion within a confined nanolitre-scale microenvironment in the AM-DMF-SCP-pro workflow. (A) Microscopy-based workflow for single-cell sample preparation in the DMF cartridge. (1) A single-cell droplet is isolated. (2) A reagent droplet is moved adjacent to the cell droplet. (3-5) The droplets are merged and mixed by sequential up-and-down movement. (6-10) Incubation. Time-stamped frames extracted from Supplementary Movie 2 illustrate the complete process, in which a single cell (highlighted by a red dashed outline) undergoes rapid lysis and digestion within 40 s and is no longer visually detectable. (B) Experimental design for digestion optimization, including two trypsin concentrations (20 and 120 ng/µL) and five digestion times (5-120 min). (C) Comparison of Missed cleavage rates across digestion conditions. (D) PGs and peptide identifications across digestion times and enzyme concentrations. (E) Median coefficients of variation (CVs) across conditions, indicating quantitative reproducibility. (F) Pairwise correlations of protein intensities across all conditions. (G) Integrated multi-parameter scoring combining digestion efficiency, proteome coverage, quantitative variability, and reaction time across conditions.

In terms of proteome coverage, both PGs and peptide identifications showed a rise-and-fall trend as a function of digestion time. At high enzyme concentration, the maximum identification depth was achieved at 15 min, with an average of 3,981 PGs and 21,258 peptides. In contrast, under low enzyme concentration, peak identification was reached at 30 min with lower values (3,554 PGs and 17,513 peptides) (Figure 4D). These results demonstrate that accelerated digestion enables higher proteome coverage within a shorter time frame. Regarding quantitative reproducibility, both enzyme conditions exhibited the lowest median CVs at 15 min (Figure 4E), indicating optimal stability at this time point. Furthermore, pairwise correlation analysis of protein expression matrices across all conditions revealed consistently high agreement (Pearson r > 0.9; Figure 4F), suggesting that variations in digestion conditions do not substantially alter global quantitative patterns.

To balance multiple performance metrics, we established a multi-criteria evaluation framework (Figure S5C), incorporating digestion time, enzyme usage, missed cleavage rate, proteome depth, and quantitative CVs. Based on their relative importance in single-cell proteomics, protein identification depth was assigned the highest weight (0.5), followed by missed cleavage rate and digestion time (0.2 each), while enzyme usage and CVs were assigned lower weights (0.05 each). After ranking and integrating these metrics, the high-enzyme, 15 min condition achieved the highest overall score, driven by its superior performance in proteome depth, digestion efficiency, and processing time (Figure 4G).

Collectively, these results demonstrate that, in nanolitre-scale reaction systems, increased enzyme concentration enables efficient 15-min digestion, achieving near-maximal proteome coverage while maintaining robust quantitative reproducibility and comparable missed-cleavage levels. This accelerated digestion improves the performance of the AM-DMF-SCP-pro workflow and increases throughput for single-cell proteomics.

### Benchmarking against plate-based workflows reveals improved signal-to-noise in nanolitre-scale SCP

To evaluate the performance of the AM-DMF-SCP-pro platform in real biological samples, we compared it with a plate-based workflow using cellenONE under identical MS acquisition conditions. HeLa single cells and blank droplets were analysed to assess proteome coverage, signal composition, and background interference (Figure 5A). The two workflows differ fundamentally in reaction architecture. AM-DMF-SCP-pro operates in ∼6 nL droplets within an oil-confined environment, minimizing exposure to air and surfaces. In contrast, the plate-based workflow uses ∼1,000 nL reaction volumes in 384-well plates and involves multiple handling steps, increasing the risk of adsorption, contamination, and dilution of effective enzyme-substrate interactions. At the protein group level, both workflows achieved comparable depth, with approximately 4,000 PGs identified per cell. However, AM-DMF-SCP-pro yielded a higher number of identified peptides, indicating improved peptide retention and enhanced sensitivity (Figure 5B). In blank droplets, the plate-based workflow produced substantially more protein group and peptide identifications, reflecting increased non-specific background arising from reagents, surfaces, and environmental contaminants. By contrast, the enclosed nanolitre-scale system effectively suppressed background accumulation (Figure 5C). Rank-intensity analysis further supported these differences. In single-cell samples (Figure 5D), both workflows detected large numbers of highly abundant intracellular proteins, including cytoskeletal and ribosomal proteins. However, the plate-based workflow also showed abnormally intense reagent-derived signals, with trypsin remaining among the top-ranked proteins, indicating substantial interference from reagent background. In contrast, these reagent-derived peaks were markedly reduced in AM-DMF-SCP-pro, and the highest-intensity proteins more closely reflected the expected distribution of abundant cellular proteins. A similar trend was observed in blank droplets (Figure 5E). In the plate-based workflow, several high-intensity entries corresponded to common contaminants in the cRAP database^41^, including keratins and trypsin, whereas both the intensity and number of these contaminant signals were substantially lower in AM-DMF-SCP-pro. A similar reduction in contaminant-derived background was evident in the raw mass spectrometric signals (Figure S6A). Representative base peak chromatograms showed characteristic trypsin-derived peaks in both blank droplets and single-cell samples processed by the well-plate workflow, whereas these peaks were markedly reduced in AM-DMF-SCP-pro and approached the noise level in single-cell samples. Consistently, overlap analysis and average-intensity comparisons of contaminant proteins further supported the lower contaminant burden in AM-DMF-SCP-pro (Figure S6B and S7). At the subcellular level, both workflows showed similar overall distributions, but AM-DMF-SCP-pro identified slightly more PGs in compartments such as the nucleoplasm and nucleolus (Figure 5F), consistent with improved sensitivity. Together, these results indicate that AM-DMF-SCP-pro improves signal-to-noise by reducing reagent-and contaminant-derived background while preserving cellular proteome depth.

**Figure 5.**
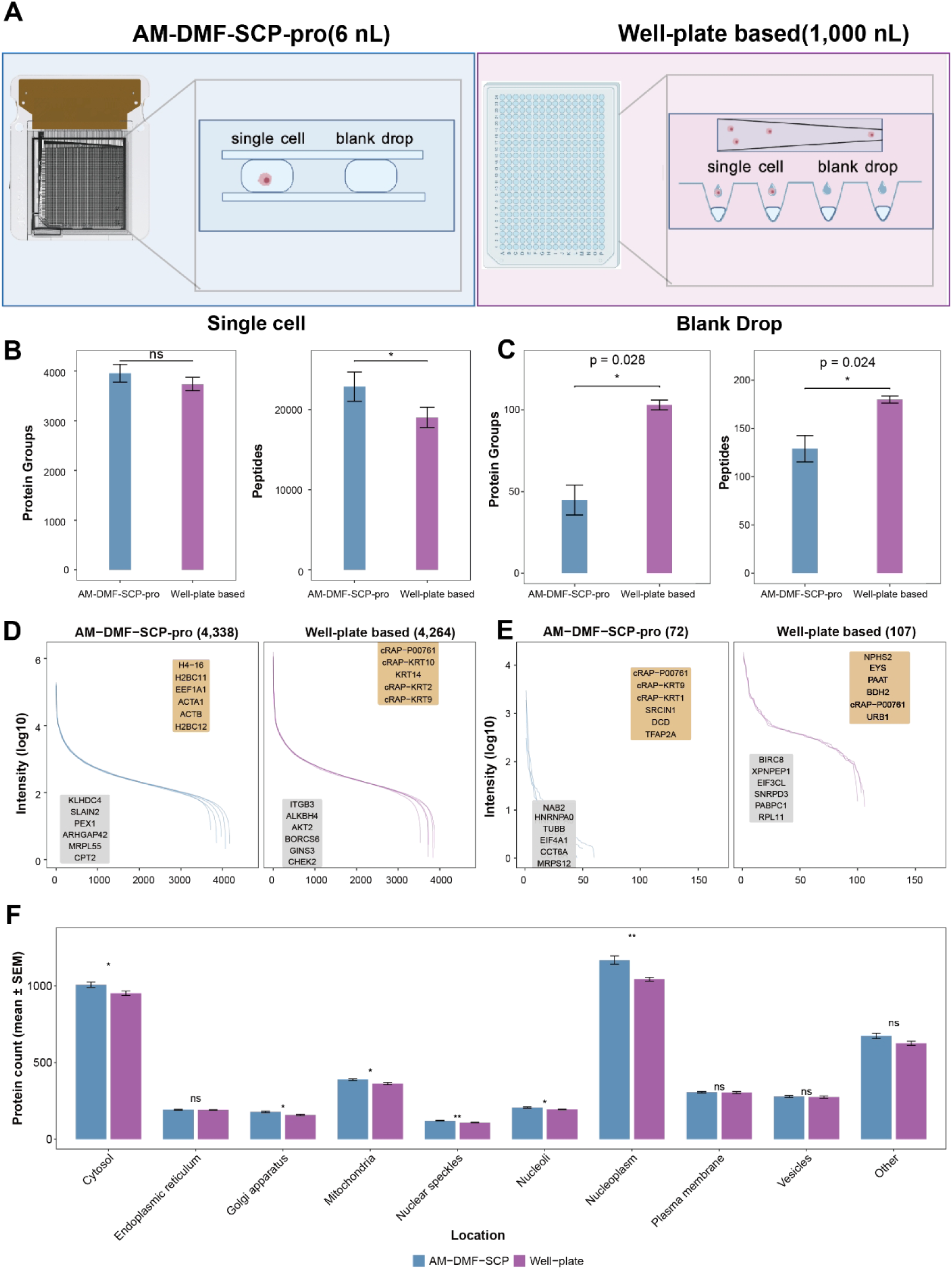
Comparison of AM-DMF-SCP-pro and well plate-based single-cell proteomics workflows. (A) Schematic comparison of AM-DMF-SCP-pro (6 nL) and a well-plate-based workflow (1000 nL). (B, C) Protein group and peptide identifications in single-cell samples (B) and blank droplets (C). Bars represent independent technical replicates, and error bars indicate s.d. (D, E) Rank-intensity distributions of quantified proteins in single-cell samples (D) and blank droplets (E), with representative proteins annotated. (F) Comparison of identified protein counts across subcellular localization categories between the two workflows.

### Single-cell proteomics resolves immune and tumor compartments in vivo

To evaluate the performance of the optimized AM-DMF-SCP-pro workflow in a complex in vivo setting, we profiled single cells from CT26 murine tumors treated with either an angiogenesis inhibitor or a vehicle control. CD45⁺ immune cells and CD45⁻ tumor cells were isolated via MACS, processed using the optimized ultrafast and streamlined nanolitre-scale single-cell proteomics workflow and analysed on an Orbitrap Astral. After stringent cell-level quality control and normalization, high-quality proteomes were retained for downstream analysis. (Figure 6A). The final dataset included 507 single cells with quantitative measurements for over 4,900 proteins, showing heterogeneous data completeness across cells (Figure 6B) and an average missing rate of ∼44%. Low-dimensional embedding by UMAP revealed a clear separation between CD45⁺ and CD45⁻ populations (Figure 6C). Consistent with immunophenotypic annotation, the pan-leukocyte marker Ptprc (CD45) was robustly detected in CD45⁺ cells and was largely absent from CD45⁻ cells (Figure 6D), supporting accurate protein-level cell identity assignment and overall data reliability. Differential protein expression analysis followed by KEGG pathway enrichment further confirmed this separation^42^: CD45⁺ cells were significantly enriched for immune-related pathways, including Fcγ receptor-mediated phagocytosis, leukocyte transendothelial migration, chemokine signalling and natural killer cell-mediated cytotoxicity, whereas CD45⁻ cells were enriched for tumor-associated programs such as DNA replication, RNA degradation, ribosome biogenesis and cell cycle regulation (Figure 6E).

**Figure 6.**
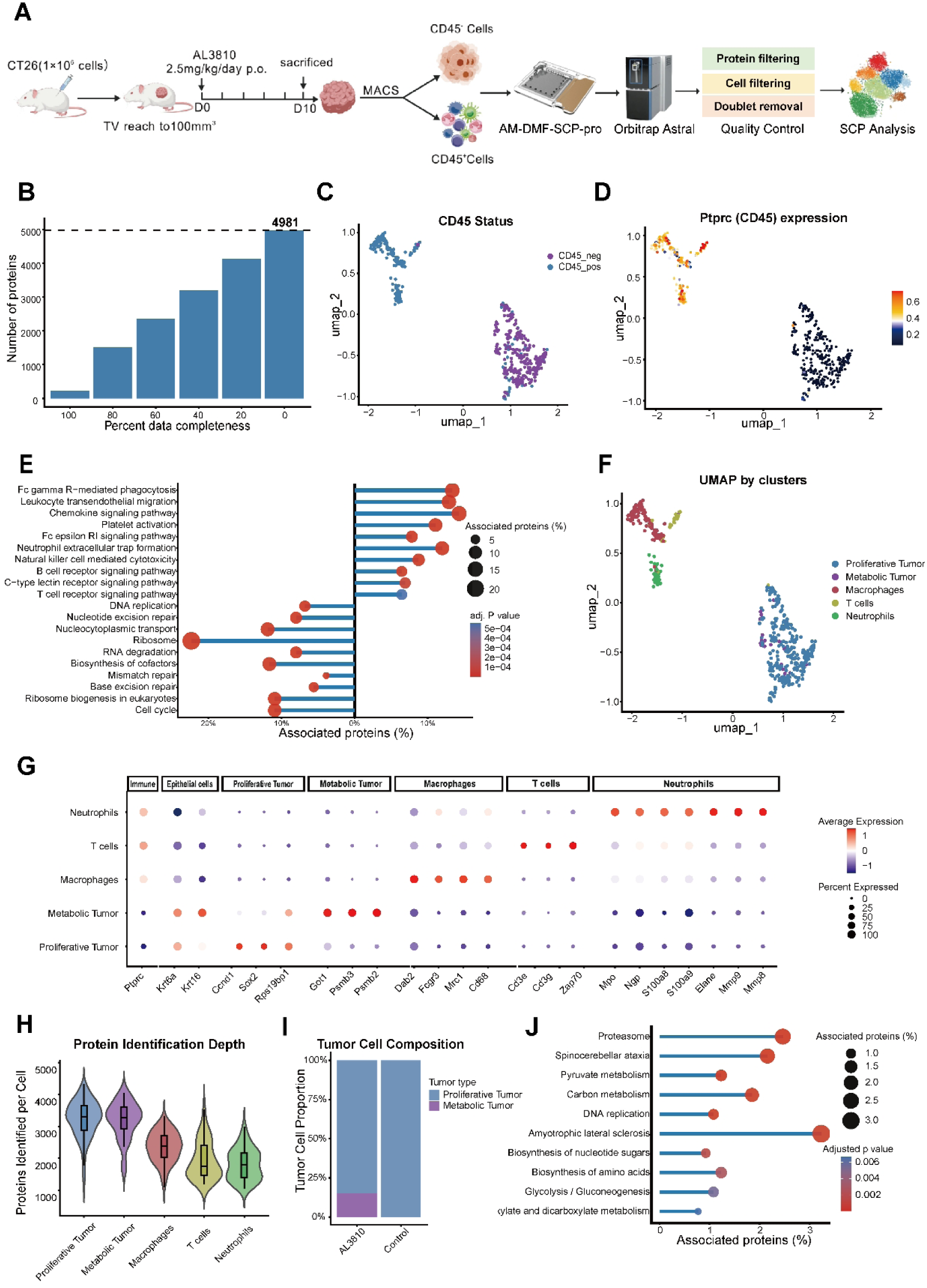
Application of AM-DMF-SCP-pro to profile cellular heterogeneity in the tumor microenvironment. (A) Workflow for CT26 tumor model establishment, treatment, tissue dissociation, MACS-based separation of CD45⁺ and CD45⁻ cell populations, AM-DMF-SCP-pro processing, and downstream quality-controlled analysis. (B) Distribution of protein identification completeness across single cells. (C) UMAP projection of single-cell proteomes colored by CD45 status. (D) UMAP projection colored by Ptprc protein expression. (E) Functional enrichment analysis of differentially expressed proteins between CD45⁺ and CD45⁻ populations. (F) UMAP projection showing clustering of major cell populations. (G) Dot plot showing the expression of representative marker proteins across annotated cell types. (H) Distribution of protein identification depth across cell types. (I) Composition of tumor cell subtypes. (J) Functional enrichment analysis of proteins associated with specific tumor cell states.

To further resolve intratumoral heterogeneity, we performed unsupervised clustering and annotated cell clusters based on canonical marker proteins. This analysis identified major cellular subtypes, including tumor cells, macrophages, T cells, and neutrophils (Figure 6F and Figure S8A). Marker protein expression patterns further validated the accuracy of cell-type annotation (Figure 6G and Figure S8B-C). A comparison of proteome coverage across cell types revealed that tumor cells exhibited the greatest depth of protein identification. In contrast, immune populations showed lower but distinct coverage profiles (Figure 6H), consistent with known differences in cell size and intracellular protein content^43^.

We next focused on the tumor cell compartment for further analysis. Two major tumor cell subtypes were identified within the tumor population: proliferative tumor cells and metabolic tumor cells. Notably, a metabolic tumor cell subpopulation was observed exclusively in the group treated with the angiogenesis inhibitor (Figure 6I). Pathway enrichment analysis of differentially expressed proteins between proliferative and metabolic tumor cells revealed significant upregulation of proteasome and glycolysis/gluconeogenesis pathways in treated tumors (Figure 6J), suggesting adaptive metabolic reprogramming of tumor cells under anti-angiogenic stress^44^.

### Single-cell proteomics enables practical neutrophil analysis in the tumor microenvironment

To complement our findings, we performed single-cell transcriptomic analysis on the same samples. However, neutrophils were not robustly detected in the transcriptomic dataset (Figure S9A-C), likely owing to their low RNA content and short transcript half-life^45^. Given the critical immunoregulatory roles of neutrophils in the tumor microenvironment^46^, we therefore performed a systematic comparison between single-cell proteomics and single-cell transcriptomics with respect to neutrophil detection and functional characterization.

Using the neutrophil granule protein database^47^, we quantified the proportion of granule proteins detected per cell, and found substantially greater granule protein coverage in single-cell proteomic data than in single-cell transcriptomic data, with the average coverage of different granule proteins categories exceeding 75% in the single-cell proteomic dataset (Figure 7A). Consistently, canonical neutrophil markers, including Ptprc, S100a8, S100a9, Ngp, Mpo, Camp, and Mmp8, ranked much higher in the proteomic dataset than in the transcriptomic dataset (Figure 7B), indicating improved detectability and signal stability of key neutrophil functional molecules at the protein level. Moreover, neutrophil granule-related signals were sparse and low in transcriptomic data but appeared more concentrated and intense in proteomic data (Figure 7C, S9D), further highlighting the advantage of single-cell proteomics for analyzing neutrophil functional states.

**Figure 7.**
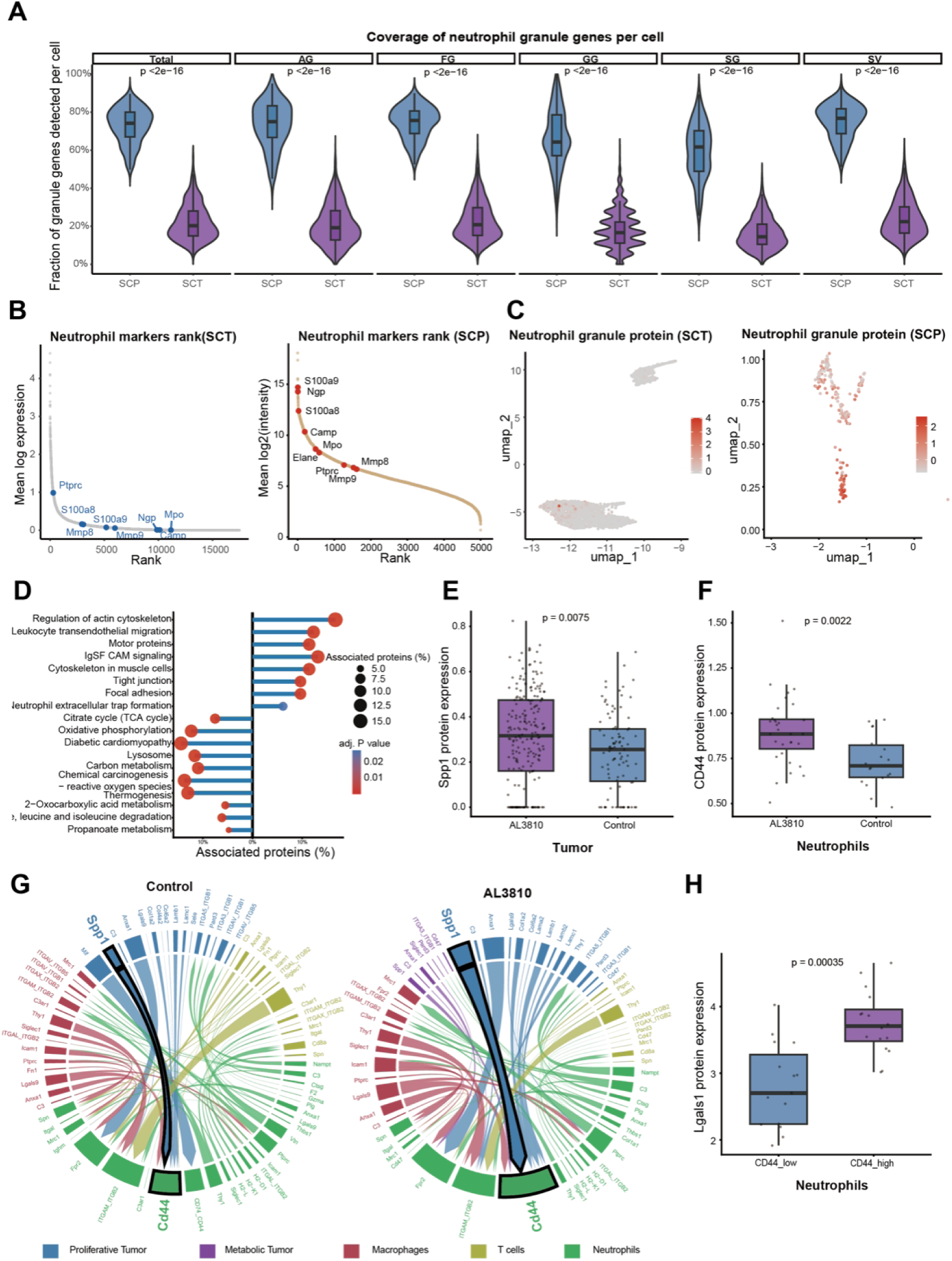
Single-cell proteomics resolves neutrophil-associated protein signatures and intercellular interactions. (A) Violin plots showing the fraction of detected neutrophil granule genes per cell across granule categories (Total, AG, FG, GG, SG, SV), comparing single-cell proteomics (SCP) and single-cell transcriptomics (SCT). (B) Rank plot of neutrophil marker genes in SCT (left) and SCP (right), with selected markers annotated. (C) UMAP projections showing neutrophil granule protein signals in SCT (left) and SCP (right). (D) Functional enrichment analysis of proteins differentially expressed in neutrophils between control (left) and treatment (right) conditions. (E) Functional enrichment analysis of proteins differentially expressed in neutrophils between control (left) and treatment (right) conditions. (F) Boxplots showing neutrophil-related signature scores across conditions in tumor cells (left) and neutrophils (right). (G) Chord diagrams illustrating ligand-receptor interactions across cell types under control (left) and treatment (right) conditions. (H) Boxplot comparing Lgals1 between CD44-low and CD44-high neutrophil populations.

Applying this framework, we examined neutrophil responses to treatment with angiogenesis inhibitors. Differential protein expression and pathway enrichment analyses revealed treatment-associated upregulation of pathways involved in cell adhesion, transendothelial migration, and neutrophil extracellular trap formation (Figure 7D). At the level of tumor-immune interactions, tumor cells exhibited increased expression of Spp1, whereas neutrophils displayed elevated expression of Cd44 following treatment (Figure 7E-F). Cell-cell interaction analysis further suggested enhanced Spp1-Cd44 signalling between tumor cells and neutrophils under angiogenesis inhibition (Figure 7G, S10A), implying that tumor cells may actively recruit Cd44⁺neutrophils by strengthening this ligand-receptor axis^48^. Consistently, flow cytometric analysis showed a higher proportion of neutrophils in the AL3810-treated group than in the control group (Figure S10B), further supporting neutrophil enrichment in the tumor microenvironment after AL3810 therapy. In addition, Cd44-high neutrophils displayed increased Lgals1 expression (Figure 7H).Given that Lgals1 is an important immunoregulatory protein that has been implicated in immune suppression, angiogenesis, and tumor progression across multiple tumor types^49^, these findings suggest that neutrophil remodeling may contribute to tumor adaptation under anti-angiogenic therapy.

Together, these results demonstrate that single-cell proteomics enables practical and robust analysis of neutrophils within the tumor microenvironment, supporting quantitative characterization of neutrophil-associated protein programs and treatment-induced changes that are challenging to capture using transcriptomic approaches alone.

## Disscussion

We developed AM-DMF-SCP-pro, a nanolitre-scale single-cell proteomics platform based on active-matrix digital microfluidics, which integrates single-cell isolation, lysis, and digestion within a single chip. Compared with conventional workflows that rely on multiple transfer steps and prolonged digestion^17,33,34^, AM-DMF-SCP-pro uses a closed, confined nanolitre-scale microenvironment together with a streamlined lysis digestion workflow to reduce total sample-preparation time to as little as 30 min, including a 15 min digestion step. This enclosed nanolitre-scale reaction system not only helps reduce the surface-related losses commonly encountered in low-input sample processing and mitigates evaporation, but also provides favorable conditions for rapid digestion by increasing effective reactant concentrations and shortening diffusion and mixing distances. Importantly, this reduction in processing time does not come at the expense of quantitative reproducibility or digestion performance, demonstrating that rapid sample preparation can be achieved in single-cell proteomics without compromising analytical quality. This workflow is further enabled by AI-guided droplet routing^37,38^, which supports precise and parallel on-chip manipulation of single-cell and reagent droplets.

Beyond accelerating sample preparation, AM-DMF-SCP-pro also improves analytical performance. Systematic optimization of MS acquisition parameters and nanolitre-scale digestion conditions increased proteome depth and detection sensitivity at the single-cell level, particularly for low-intensity, low-abundance proteins. Compared to well-plate-based workflows, the oil-confined droplet system substantially reduced reagent and environment derived background, improving signal-to-noise while preserving cellular proteome depth. This is especially important for trace single-cell analysis, in which background accumulation can directly determine whether low-abundance proteins and weak signalling pathways are reliably detected^7,9,10^. Meanwhile, comparable CV distributions, cross-workflow correlations, and missed-cleavage levels indicate that these gains are achieved without compromising quantitative performance.

Beyond its methodological advances, AM-DMF-SCP-pro also performed well in complex biological samples. In murine tumors, the platform reliably resolved immune and tumor compartments in vivo and further identified treatment-associated tumor cell states. In the angiogenesis inhibitor-treated group, we observed a distinct metabolic tumor cell subpopulation together with enrichment of proteasome and glycolysis/gluconeogenesis pathways, suggesting adaptive metabolic reprogramming under therapeutic stress. The platform also enabled quantitative single-cell proteomic analysis of neutrophils in the tumor microenvironment, revealing neutrophil-associated protein programs and treatment-induced changes. Combined with single-cell transcriptomic analysis, cell-cell communication analysis, and flow cytometry validation, these findings highlight the added value of single-cell proteomics for studying functional cell states in complex tissues.

Although this study demonstrates the potential of AM-DMF-SCP-pro for single-cell proteomic sample preparation and applications in complex tissues, several aspects still warrant further improvement. First, although the overall workflow time has been substantially reduced, the sample throughput of a single chip remains limited. Given that the active-matrix digital microfluidic chip is built on a mature thin-film semiconductor manufacturing process from consumer electronics, and in light of our ongoing efforts to expand chip size and pixel scale, we anticipate that the throughput of a single chip can be further increased. At the same time, the overall analytical throughput is still constrained by mass spectrometry acquisition capacity, and further improvements in platform performance will depend on continued advances in acquisition speed and detection efficiency. Second, although the platform has been validated here using HeLa cells and murine tumor samples, its robustness and generalizability across a broader range of primary cell types, clinical samples, and more complex tissues remain to be established. Overall, AM-DMF-SCP-pro not only provides a new technical route for rapid, low-background, and scalable single-cell proteomic sample preparation, but also offers a foundation for resolving functional cell states and treatment-associated remodeling in complex tumor microenvironments at proteome level.

## Methods

### Cell Culture and harvested

HEK-293T and HeLa cells were cultured in DMEM supplemented with 10% FBS (Gibco) and 1% penicillin-streptomycin (Sigma-Aldrich) at 37°C under 5% CO₂. Cells were harvested at approximately 80% confluency through 1-minute trypsinization at 37°C, quenched with complete medium, pelleted at 1,000 g for 1 minute, and washed three times with ice-cold PBS before proceeding to bulk or single-cell proteomic analyses. Escherichia coli W3110 was grown overnight in M9 minimal medium at 37°C with shaking (220 rpm), collected by centrifugation at 2,000 g for 5 minutes at 4°C, and washed twice with ice-cold PBS. Saccharomyces cerevisiae was revived in YPD medium at 30°C with shaking (222 rpm), expanded into fresh YPD for 24 hours, harvested by centrifugation, and washed twice with PBS prior to standard preparation.

### Animal studies

Female Balb/c mice aged 6-8 weeks old were purchased from Shanghai Jihui Animal Breeding Laboratory and were housed under specific pathogen-free conditions. All animal experiments were conducted following the approval of the Institutional Animal Care and Use Committee (IACUC) of the Shanghai Institute of Materia Medica, Chinese Academy of Sciences. CT26 cells were subcutaneously inoculated into the right side of the flank at 1×10^6^ cells per implant. Animals were randomized to receive oral administration of vehicle or a VEGFR inhibitor when the tumor volume reached about 100 mm^3^.

### Tumor Harvesting and Magnetic separation of CD45+ and CD45-cells

Tumor-bearing mice were sacrificed after drug treatment for 10 days. CT26 tissues were cut into small pieces and digested with Collagenase IV (Worthington, 1 mg/mL) and DNase I (Roche, 0.1 mg/mL) at 37 °C for 45 min with gentle agitation. The dissociated cells were filtered through a 70 μM cell strainer. Erythrocytes were lysed with Red Blood Cell Lysis Buffer (Meilunbio) for 3 min, and cells were washed with PBS. After that, cells were counted and centrifuged at 300g for 5 min. The cell pellet was resuspended in MACS buffer (PBS containing 0.5% bovine serum albumin (BSA) and 2 mM EDTA) at a density of 1×10^7^ cells per 90 μL and incubated with CD45 TIL MicroBeads (Miltenyi Biotec, 10 μL per 1×10^7^ cells) for 15 min at 4 °C. The suspension volume was adjusted to 500 μL with MACS buffer, and magnetic separation was performed using LS columns placed in a MACS Separator according to the manufacturer’s instructions. The flow-through fraction, containing CD45-cells, was collected, and magnetically retained CD45+ cells were subsequently eluted after removal of the column from the magnetic field. Both fractions were collected and used for downstream analyses.

### Design and fabrication of the DMF cartridge

Each DMF cartridge consisted of, from top to bottom, an injection-molded plastic housing, an indium tin oxide (ITO)-coated glass top plate (thickness 0.5 mm), an adhesive gap spacer, and a bottom plate integrating a bonded flexible printed-circuit (FPC) connector (Figure 1F). The bottom plate was a thin-film-transistor (TFT) backplane fabricated by a standard amorphous-silicon (a-Si) TFT process. The uppermost layer of the TFT fabrication process was a patterned ITO interdigitated driving electrode array comprising 128 × 128 individually addressable, approximately square electrodes (250 µm × 250 µm each). A ∼300 nm silicon-nitride dielectric was deposited over the electrodes, followed by spin-coating of a ∼50 nm hydrophobic layer of Teflon-AF 1600X (Chemours).

The top plate was ITO glass (thickness 0.5 mm) and was laser-drilled with 1 mm and 3 mm diameter access holes serving as sample inlet and outlet, respectively, The ITO surface was likewise spin-coated with Teflon-AF 1600X. The top and bottom plates were bonded using an adhesive gap spacer comprising a UV-curable adhesive premixed with 50 µm spacer beads, defining a uniform ∼50 µm inter-plate gap. Electrical connection was established by thermocompression (hot-press) bonding of the FPC to the bottom plate. The injection-moulded plastic housing was bonded to the top plate using an epoxy adhesive to ensure sealing during sample loading, and each cartridge carried a QR code label for traceability.

### Control System and Operation of the DMF Cartridge

The DMF cartridge was operated within the BOXmini-SCP platform for single-cell proteomic sample preparation. The cartridge connects to the instrument via an FPC connector at its base, enabling independent addressing of all on-chip electrodes for precise droplet actuation. Platform control is implemented through dedicated software that integrates electrical driving, dual-scale imaging, motorized positioning, and temperature regulation. The system incorporates a high-speed electronic driving module for independent addressing of the TFT array and driving electrodes to perform droplet generation, merging, mixing, dispensing, and fully automated single-cell sample preparation. A dual-scale optical imaging module provides a 280× high-magnification path for single-cell identification, localization, and selection with a built-in light source, and an 8× low-magnification path for real-time monitoring of global droplet operations using ring illumination to enhance droplet-edge contrast. Both optical paths are mounted on a triaxial motorized positioning stage that enables precise X/Y/Z movement for rapid targeting of cells and droplets, as well as dynamic calibration of operating sites. For on-chip biochemical reactions, the cartridge is mounted on an integrated thermoelectric cooler (TEC, Peltier module) that provides a tunable 0-100 °C temperature range for accurate control of critical steps, such as cell lysis and enzymatic digestion.

### Automated on-chip single-cell sample preparation

The detailed AM-DMF-SCP workflow has been described previously. For AM-DMF-SCP-pro, the DMF cartridge was first primed by filling the entire chamber with 50 µL of immiscible silicone oil (PMX-200, 2 cSt, Dow). After inserting the cartridge into the BOXmini-SCP platform and initializing the control software, approximately 1 µL of cell suspension (3.5 × 10^5^ cells/mL) and 0.3 µL of premixed lysis/digestion buffer (0.4% DDM, 40 ng/µL trypsin, 200 mM TEAB) were dispensed into designated on-chip reservoirs. Under automated DMF control, the cell suspension was discretized into 320 droplets and the premix into 96 droplets. Each cell droplet was sequentially imaged using the high-magnification optical path; droplets containing exactly one cell were selected for processing, while empty or multicell droplets were routed to the on-chip waste region. All premixed reagent droplets were then routed to the selected cell droplets, which were merged and mixed in parallel to initiate lysis and digestion. The integrated thermoelectric cooler was automatically set to 50 °C for incubation, after which the temperature was returned to ambient. The single-cell digests were transferred to the outlet port and collected into a 384-well plate, ready for either immediate LC-MS/MS analysis or storage at −80 °C.

### Enhanced SIPP-based multi-droplet routing

For automated multi-droplet routing on the digital microfluidic platform, we implemented an enhanced safe-interval path planning (enhanced SIPP) framework by extending our earlier SIPP-based routing approach. Droplets were first ordered according to a predefined spatial priority, and candidate destination sites were filtered to exclude invalid or inaccessible positions before goal assignment. Each droplet was then planned sequentially on the chip grid under both static constraints imposed by fixed device structures and dynamic spatiotemporal constraints introduced by previously planned droplets. The enhanced framework explicitly accounts for droplet footprint, safety buffer, time-indexed occupancy and final-position holding, with optional edge-conflict constraints to prevent swap conflicts between neighbouring droplets. For recovery operations, droplets could be routed to a designated waste region rather than to a single target position. Residual conflicts, when present, were resolved by inserting waiting steps into lower-priority paths. The enhanced SIPP workflow, including priority-ordered planning, spatiotemporal reservations and optional post-processing, is summarized in Supplementary Algorithm 1. Planning time and makespan were evaluated in benchmark scenarios containing 10-100 droplets and compared with those obtained using the previous SIPP-based implementation.

### NanoLC-MS/MS analysis

Single-cell proteomic analyses were performed on an Orbitrap Astral mass spectrometer (Thermo Fisher Scientific) coupled to a Vanquish Neo UHPLC (Thermo Fisher Scientific) and operated with a FAIMS Pro Duo interface (Thermo Fisher Scientific). The electrospray voltage was set to 1,900 V in positive ion mode. We used Aurora Elite TS analytical columns (15 cm × 75 μM, IonOpticks) with a nanoflex ion source, a homemade oven for 50 °C, and a homemade column adaptor. FAIMS was operated in standard-resolution mode with a compensation voltage. Mobile phase A and B were 0.1% formic acid in water and 0.1% formic acid in 80% acetonitrile, respectively. Samples were injected with direct-injection mode, and separation was carried out under a 12-min gradient at 600 nL/min unless otherwise noted. The gradient program was as follows: 4% B for 0-1.0 min, increased to 8% B at 1.0 min, 28% B at 9.0 min, and 40% B at 10.2 min. The column was then washed by ramping to 99% B at 10.6 min and held until 12 min. During washing, the flow rate was increased to 700 nL/min between 11.0 and 12 min to enhance column flushing. For DIA acquisition, MS1 spectra were acquired in the Orbitrap at a resolution of 240,000 over m/z 400-800, with a custom AGC target of 5,000,000 and a maximum injection time of 100 ms. MS2 spectra were acquired in the Astral analyser covering the precursor mass range m/z 400-800. Fragmentation was performed by HCD using a normalized collision energy (NCE) of 25%. MS2 spectra were collected over m/z 100-1700 in centroid mode and an AGC target set to 800% (absolute AGC target 80,000). The DIA loop control was time-based with a cycle time of 0.6 s.

### Method optimization for ultra-low input data acquisition

To identify acquisition settings suitable for ultra-low-input samples, DIA methods were systematically evaluated by varying the DIA isolation window width (Th) and the FAIMS compensation voltage (CV) while keeping chromatography and other MS parameters constant. Candidate methods were compared using proteome coverage (PGs and peptides) and quantitative reproducibility (coefficient of variation across technical replicates). The final acquisition method used for all subsequent single-cell analyses employed a 20-Th isolation window and a FAIMS CV at −48 V, as th**e**se settings provided the best overall balance across the evaluation criteria.

### Quantitative performance assessment using mixed-species peptide standards

To evaluate quantitative accuracy and ratio fidelity of the optimized workflow under ultra-low-input conditions, mixed-species peptide standards were prepared using tryptic digests from Escherichiacoli (E), Homosapiens (H), and Saccharomycescerevisiae(S). Peptides from each species were digested and quantified independently, then combined at predefined mass ratios to generate mixtures with known inter-species fold differences. Three ratio schemes were designed to assess quantitative performance across moderate and extended dynamic ranges, including E:H:S = 2:5:3 and 3:5:2, E:H:S = 1:5:4 and 4:5:1, and E:H:S = 1:10:9 and 9:10:1. All mixtures were diluted to a total peptide input of 50 pg per injection, matching the peptide amounts used in single-cell proteomics experiments. Protein-level intensities were used to calculate measured log₂ fold changes between species, which were compared with theoretical ratios defined by the mixing design to assess quantitative accuracy, intensity dependence, and reproducibility across the tested ratio schemes.

### MS raw data analysis

For the analysis using Spectronaut v.19.3 (Biognosys), raw files were processed using a library-free directDIA+ approach or method evaluation mode, employing with a library-free search against the Homo sapiens Swissprot reference proteomes database (release 2021_09, 20588 entries) or Musmusculus(release 2024_08, 17207 entries) plus the common Repository of Adventitious Proteins (cRAP) database(125 entries). In the analysis, cross-run normalization was closed.

### Single cell proteomics cell filtering

Cells with fewer than 20% of the maximum number of identified proteins in the dataset were excluded. Proteins detected in at least 1% of cells were retained, and missing values were treated as undetected signals and set to zero. Potential doublets were identified and removed using scDblFinder^50^ (v1.22.0), and the remaining cells were used for downstream analyses.

### Single cell proteomics Seurat pipeline

The single-cell proteomics data were analysed using Seurat^51^ (v5.3.1). The intensity data were normalized as described above, and this normalized matrix was used as input for Seurat. FindVariableFeatures() was applied with nfeatures = 2000 to select highly variable proteins. Principal component analysis was performed with the total number of principal components set to 28. The data were batch-corrected using Harmony^52^ (v1.2.4), with processing date used as the grouping variable. FindNeighbors() was run on the Harmony-corrected embeddings using 28 dimensions, and FindClusters() was performed using the Leiden algorithm. The number of clusters was determined based on the ElbowPlot() method and cluster stability across multiple resolutions, and was ultimately set to 6.

### Single cell population markers

For the population markers the FindMarkers() function in Seurat was used. Differential analysis was performed using the Wilcoxon rank-sum test. Proteins detected in at least 25% of cells and with a log2 fold change greater than 0.5 were considered candidate markers. P values were adjusted using the Benjamini-Hochberg method, and adjusted p < 0.05 was considered statistically significant.

### Cell-cell communication analysis

Cell-cell communication analysis was performed using CellChat^53^ (v1.6.1). CellChatDB. mouse was used as the ligand-receptor reference database. Only signalling-related proteins annotated in the database were retained. Communication probabilities were inferred using the truncated mean method with trim = 0.1, and probability estimation used 100 bootstrap iterations. Interactions involving cell groups with fewer than three cells were excluded. Pathway-level communication probabilities were aggregated to generate the global communication network for downstream visualization.

### Copy number calculation

Protein copy numbers were estimated using the proteomic ruler^54^, which uses histone abundance as an internal reference to infer absolute protein copy numbers per cell. Copy number estimation was performed based on the relative iBAQ-like intensity matrix.

### Single-cell transcriptomics analysis pipeline

Single-cell transcriptomics data were imported using Read10X and processed with Seurat (v5.2.1). Cells with fewer than 500 or more than 6,000 detected genes, total UMI counts greater than 40,000, mitochondrial gene content greater than 10%, or hemoglobin gene content greater than 5% were excluded.

Genes detected in fewer than three cells were removed. Potential doublets were identified and removed using scDblFinder (v1.14.0), and ambient RNA contamination was corrected using decontX (v1.4.0). Data normalization was performed using SCTransform. The top 3,000 highly variable genes were used for PCA, and the first 29 principal components were used to construct the nearest-neighbor graph and perform Leiden clustering.

### Data visualization

Dot plots, UMAP plots, feature plots, and heatmaps were generated using Seurat (v5.3.1). Violin plots, box plots, scatter plots, and bar plots were generated using ggplot2 (v4.0.1). Chord diagrams for cell-cell communication were generated using circlize (v0.4.16), and communication bubble plots were generated using ggplot2 (v4.0.1).

## Data and code availability

Single-cell proteomics raw data and search file results have been deposited to the ProteomeXchange Consortium via the iProX^55,56^ partner repository with the dataset identifier PXD077501 and PXD073385. The code for data analysis and plot is available at https://github.com/ youngbee12/AM-DMF-SCP-pro. https://www.iprox.cn/page/PSV023.html;?url=1777439561143bpi2 https://www.iprox.cn/page/PSV023.html;?url=1777439918607og8R

## Additional information

Supporting Information: Supporting figures, Algorithm, and Movies.

## Author Contributions

K.J. and Z.Y. conceived and designed the study. K.J., Z.Y., A.D. and R.W. performed the experiments, acquired and analysed the data, and wrote the original draft. Y.W. provided experimental assistance. Y.M., Y.W., M.T. and Q.L. provided guidance on data analysis. J.J., C.C. and S.H. contributed to software algorithm development and support. W.D., Y.C., H.M. and H.Z. conceived and supervised the work. All authors contributed to revising the manuscript and approved the final version.

## Acknowledgment

This work was financially supported by the National Key Research and Development Program of China (2025YFA1309400, 2023YFF0721500), the Innovative Drug Research and Development National Science and Technology Major Project (2025ZD1804100), the National Natural Science Foundation of China(22425703), the Shanghai Leading Talent Program of Eastern Talent Plan (LJ2023123 to H.Z.), the Strategic Priority Research Program of the Chinese Academy of Sciences (XDB0830000 to H.Z.), the Natural Science Foundation of Shandong Province (ZR2024QC161), the Suzhou Basic Research Pilot Project (SSD2023013, SSD2024023), Guangdong Scientific and Technological Project (2025A0505010003).

