## Supplementary material for "AM-DMF-SCP-pro: An Ultrafast and Streamlined Nanolitre-Scale Lab-on-a-Chip Platform for Single-Cell Proteomics and Tumor Microenvironment Profiling": FIgure S

### Supplementary Figures

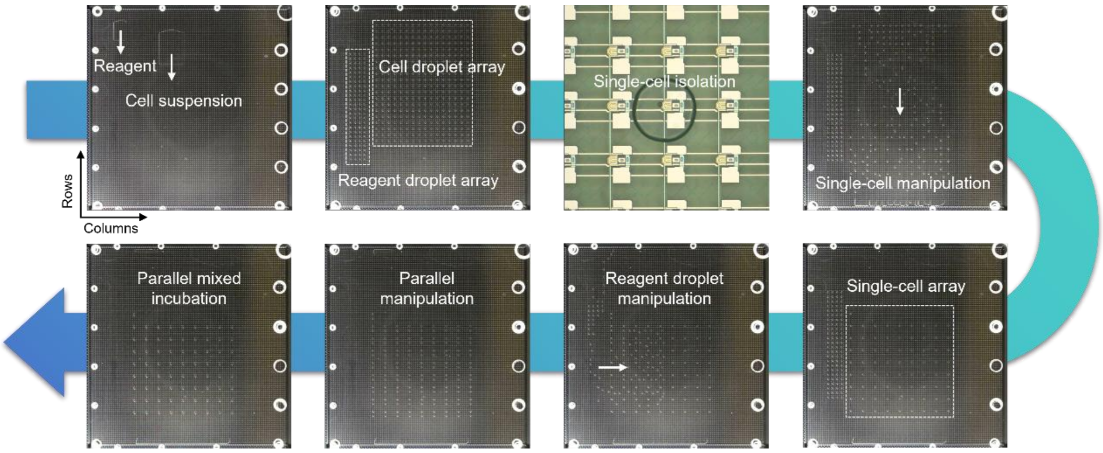


**Figure S1 Photographs corresponding to the schematic workflow in Figure 2E.** Representative photographs illustrate key stages of single-cell sample preparation on the DMF cartridge, including sample loading, droplet splitting into arrays, single-cell droplet isolation, droplet routing, droplet merging and mixing, and on-chip incubation.

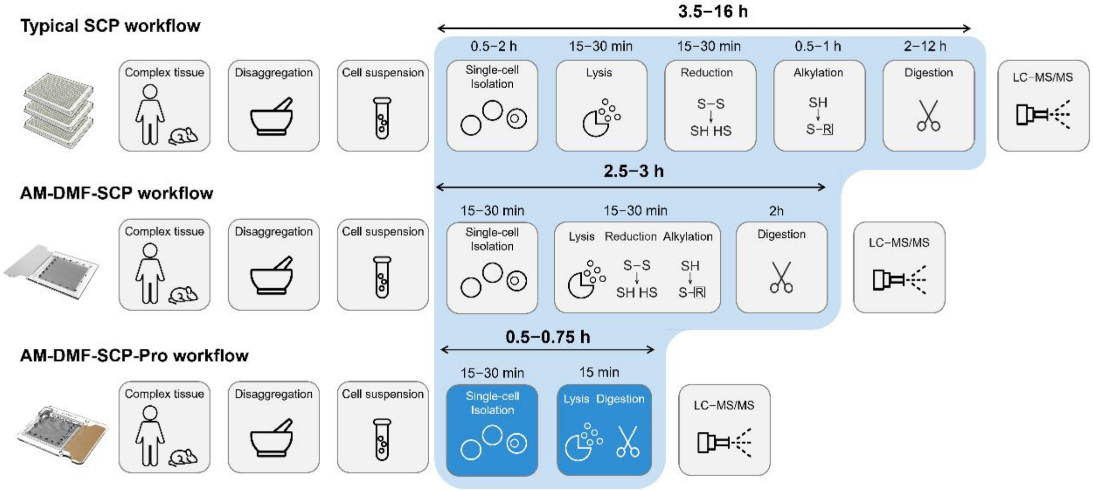


**Figure S2 Comparison of the proposed ultrafast, automated lab-on-a-chip SCP workflow with AM-DMF-SCP and a conventional SCP workflow.** Comparison of workflow design, reaction steps, and total sample-preparation time among AM-DMF-SCP-pro, AM-DMF-SCP, and a conventional SCP workflow.

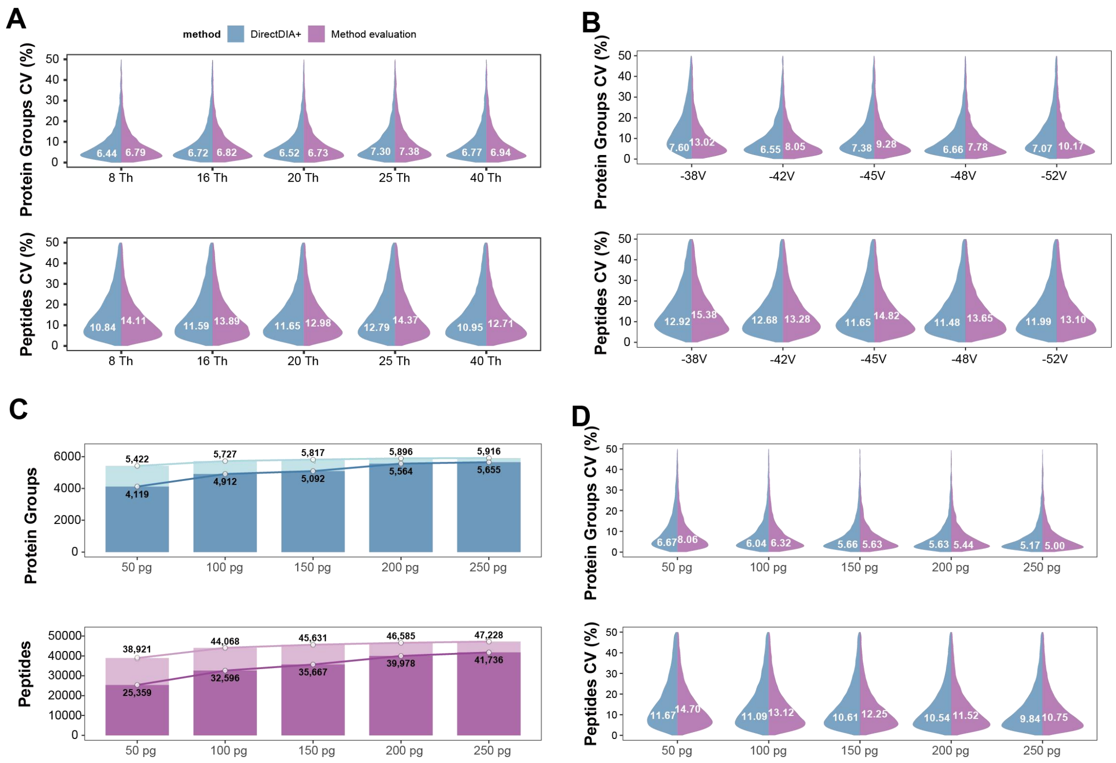


**Figure S3 Effects of DIA window width, FAIMS CV, and sample loading on identification and variation**

(A) CV distributions under different DIA window widths.

(B) CV distributions under different FAIMS compensation voltages.

(C) Protein-group and peptide identifications at different sample loading amounts.

(D) CV distributions at different sample loading amounts.

E. coli : H. sapiens : S. cerevisiae

2 : 5 : 3 / 3 : 5 : 2

| | | |
|---|---|---|
| E. coli | 956 | 1061 |
| H. sapiens | 3243 | 3283 |
| S. cerevisiae | 1371 | 1226 |
| Total | 5570 | 5570 |

E. coli : H. sapiens : S. cerevisiae

1 : 5 : 4 / 4 : 5 : 1

| | | |
|---|---|---|
| E. coli | 824 | 1219 |
| H. sapiens | 3398 | 3417 |
| S. cerevisiae | 1492 | 1245 |
| Total | 5714 | 5881 |

E. coli : H. sapiens : S. cerevisiae

1 : 10 : 9 / 9: 10 : 1

| | | |
|---|---|---|
| E. coli | 851 | 1247 |
| H. sapiens | 3787 | 3789 |
| S. cerevisiae | 1670 | 1040 |
| Total | 6308 | 6076 |

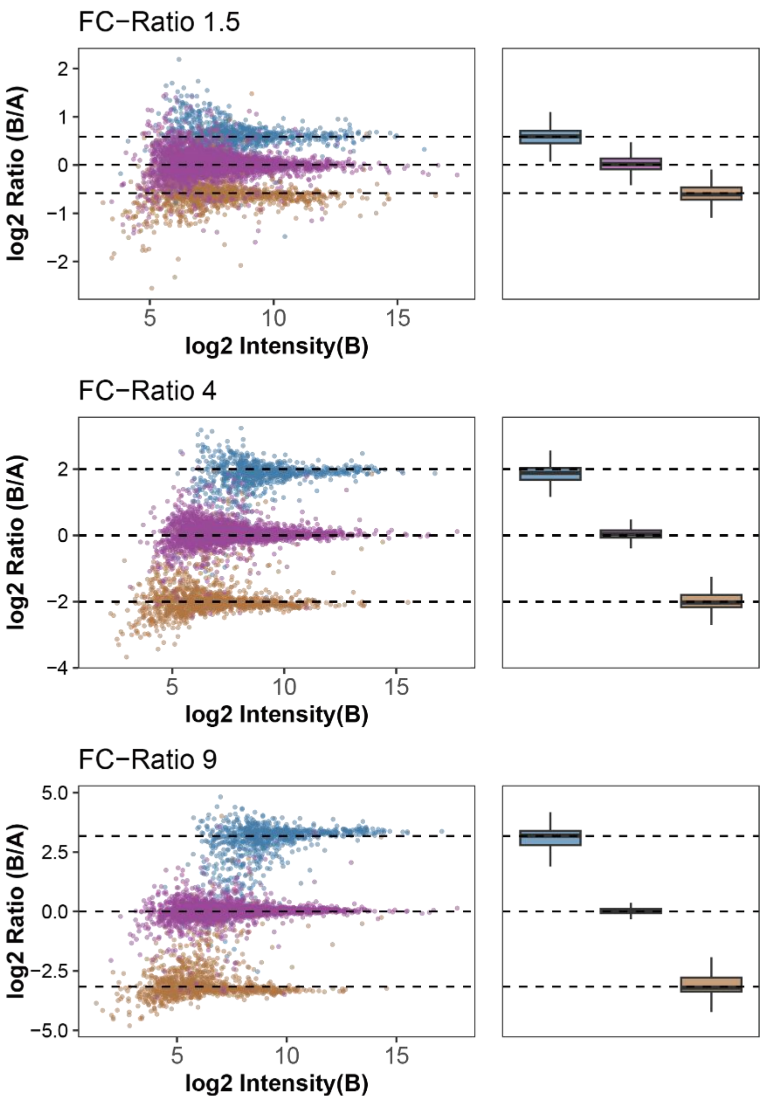


**Figure S4 Quantitative Performance Evaluation Using a Three-Species Proteome Mixture**

Quantitative assessment of the AM-DMF-SCP-pro workflow using mixed proteomes from E. coli, H. sapiens, and S. cerevisiae at different predefined fold-change ratios. Left, numbers of quantified proteins from each species in the two comparison groups. Middle, MA plots showing log2 ratios versus protein intensity for proteins from the three species. Right, box plots showing the distribution of protein ratios for each species.

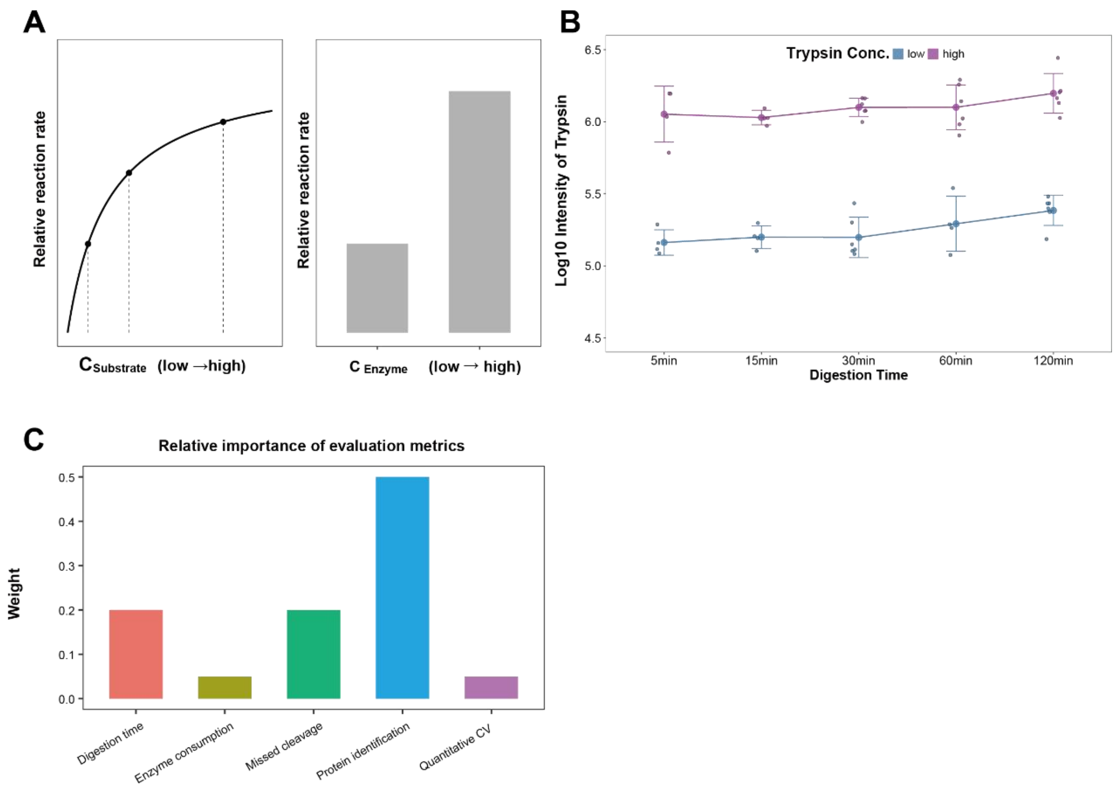


**Figure S5 Conceptual Framework and Evaluation Metrics for Optimization of On-Chip Digestion Conditions**

(A) Conceptual relationship between enzymatic reaction rate and substrate/enzyme concentration under nanolitre-scale conditions.

(B) Relative intensity of trypsin under different concentration settings.

(C) Predefined weighting of five evaluation metrics used for optimization of single-cell proteomics experiments.

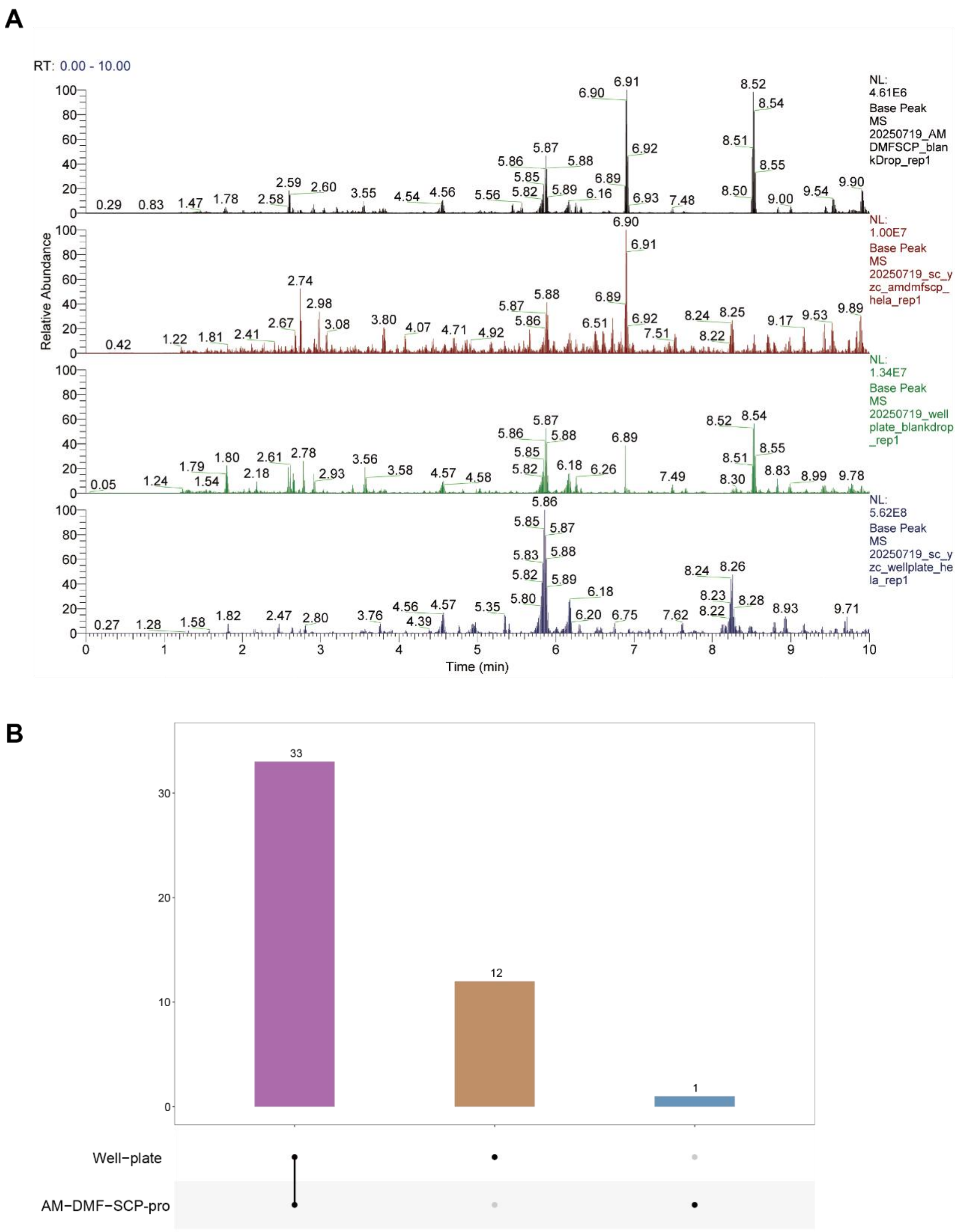


**Figure S6 Comparison of Background Signals and Contaminant Proteins Between AM-DMF-SCP-pro and Well-Plate Workflows**

(A) Representative base peak chromatograms (BPCs) of blank droplets and single-cell samples processed by the AM-DMF-SCP-pro and well-plate workflows. From top to bottom: AM-DMF-SCP-pro blank droplet,

AM-DMF-SCP-pro single cell, well-plate blank droplet, and well-plate single cell.

(B) UpSet plot showing the numbers and overlap of contaminant proteins identified in single-cell samples processed by the AM-DMF-SCP-pro and well-plate workflows.

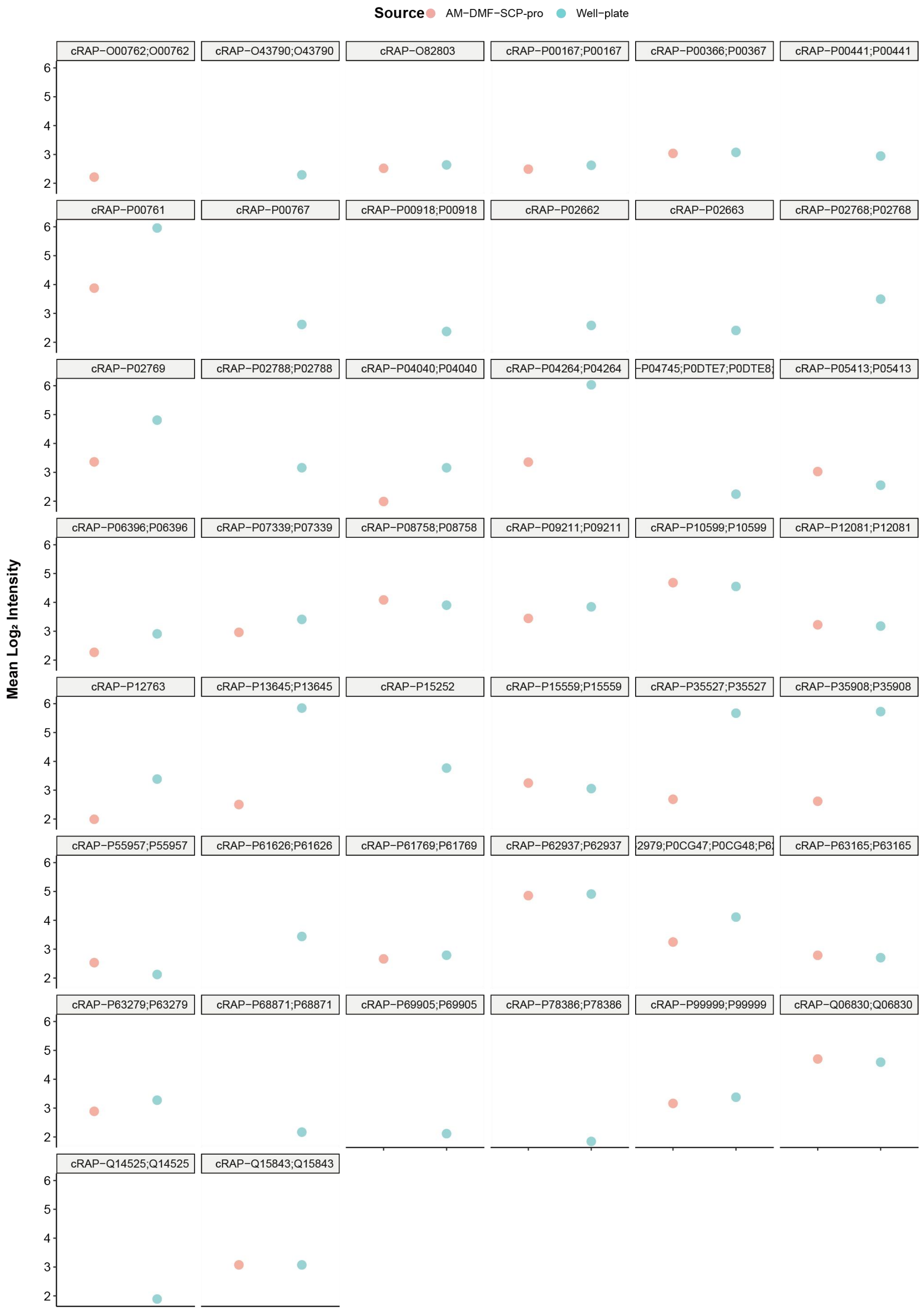


**Figure S7 Average Intensities of Contaminant Proteins Identified in both Workflows**

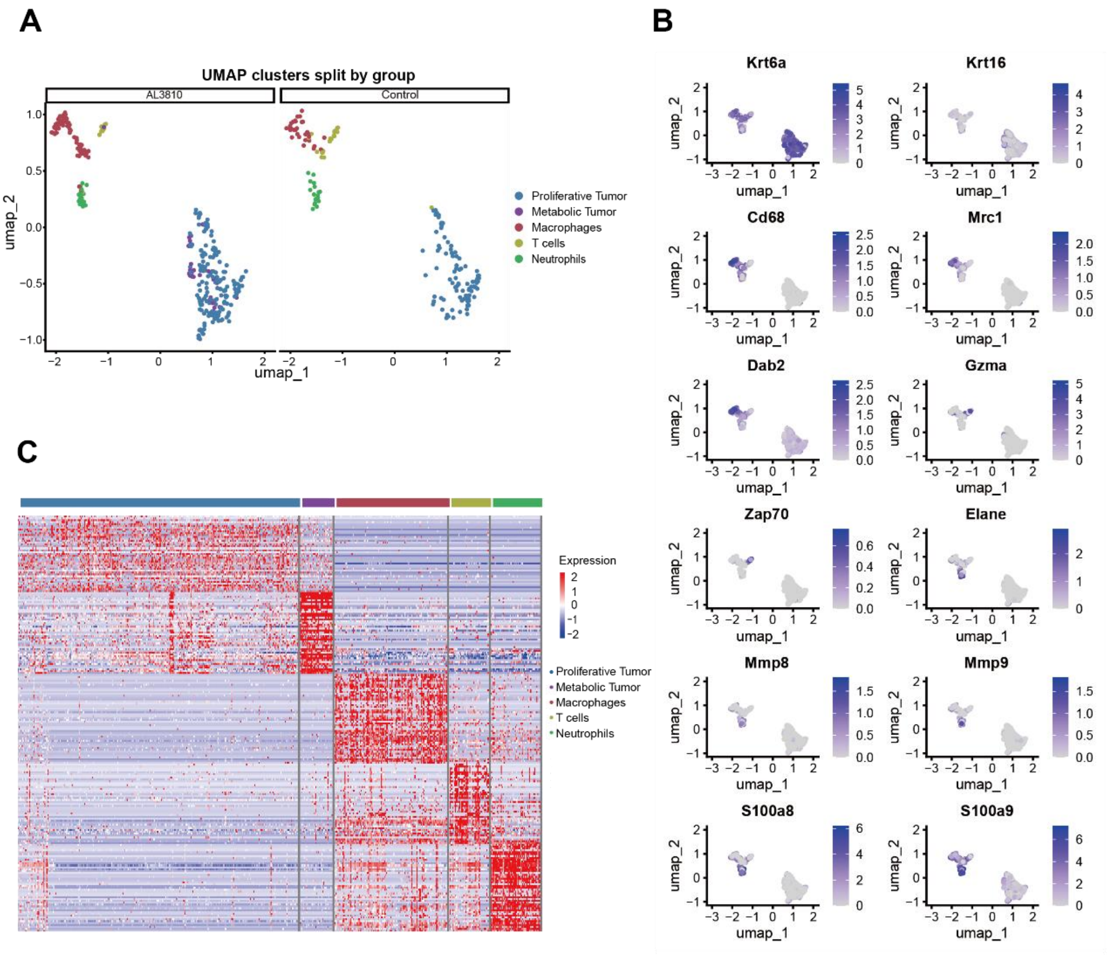


**Figure S8 Single-cell proteomic clustering and marker protein expression in tumor microenvironment samples**

(A) UMAP projection of single-cell proteomic data displayed by experimental group.

(B) Expression distribution of representative marker proteins in the UMAP space.

(C) Heatmap showing protein expression patterns across annotated cell types.

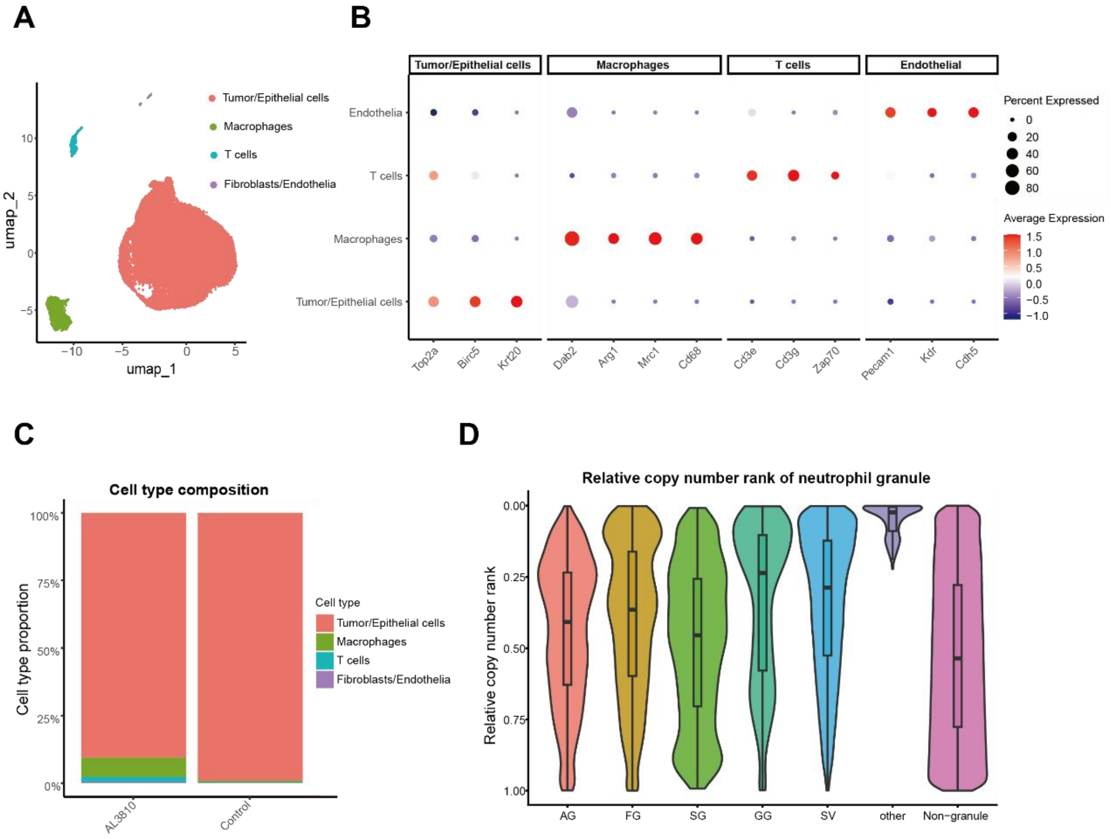


**Figure S9 Cell-type annotation from single-cell transcriptomics and neutrophil granule protein abundance in single-cell proteomics**

(A) UMAP visualization of single-cell transcriptomic clusters.

(B) Dot plot showing the expression of representative marker genes across annotated cell types.

(C) Relative proportions of annotated cell types in each sample group.

(D) Violin plot showing the relative copy-number ranks of neutrophil granule proteins in single-cell proteomic data.

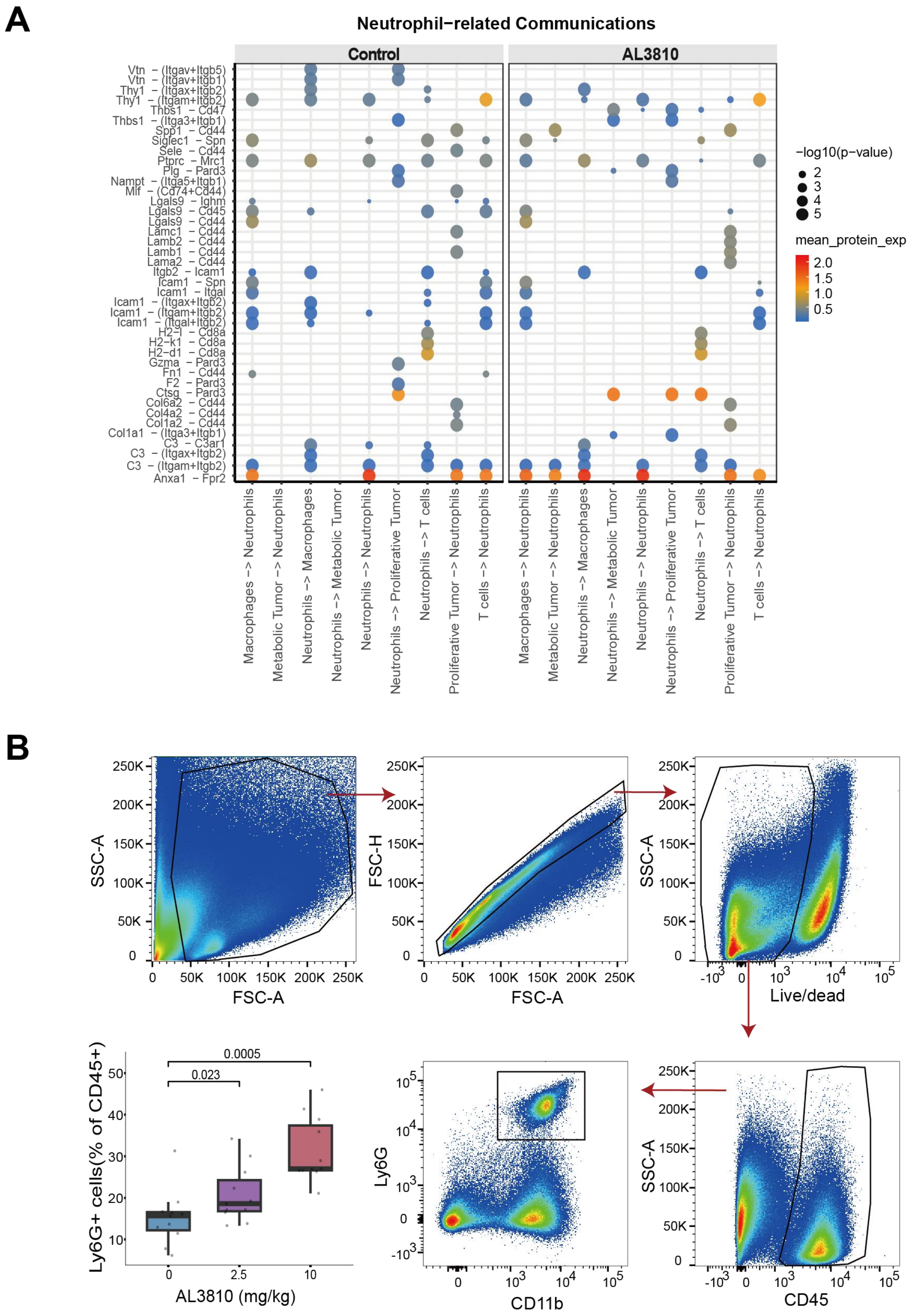


**Figure S10 Neutrophil-associated cell communication analysis and flow-cytometric validation of neutrophil abundance**

(A) Bubble plot showing inferred neutrophil-related cell–cell communication networks in the Control and AL3810 groups.

(B) Flow-cytometry gating strategy and quantification of neutrophil proportions within immune cells under different treatment conditions.

### Algorithm S1. Enhanced safe-interval path planning for multi-droplet routing

Input: grid, fixed_obstacles, droplets, candidate ends, drop_area
Output: time-indexed paths for all droplets

1. Initialize map and reservation table R.
2. Sort droplets by priority.
3. Filter and sort candidate end positions.
4. Assign goals to non-recovery droplets with conflict checking.
5. For each droplet in priority order:
    a. Set goal = drop_area for recovery droplets, otherwise use assigned point goal.
    b. Plan a path using the enhanced SIPP planner under static obstacles and dynamic reservations.
    c. Store the path and update vertex, edge, and goal-hold reservations.
6. If residual conflicts remain, resolve them by priority-based wait insertion.
7. Return all paths.

Enhanced SIPP planner:
- State: (node_position, safe_interval, t_arrive)
- Safe intervals consider inflated static obstacles and dynamic reservations
- Heuristic:
    POINT: Manhattan distance to point goal
    AREA: minimum Manhattan distance to goal-region boundary

- Successors: 4-neighborhood moves with implicit waiting
- Goal test:
  POINT: exact goal position
  AREA: droplet rectangle fully inside goal region

Conflict resolver:
- Process droplets in priority order
- Build reservations from earlier paths
- Insert wait steps at the start until conflicts are removed or a safety limit is reached
